# Detection of Stress in Naturalistic Settings Through Passive Mobile Sensing

**DOI:** 10.64898/2026.08.28.745906

**Authors:** Selin Acan, Peter De Looff, Mitra Baratchi, Martin Dresler, Erno Hermans

## Abstract

Unobtrusive stress detection using wearable sensors could enable scalable, continuous mental-health monitoring. However, stress is an inherently subjective state that can only be inferred indirectly from physiological signals, making generalizable detection in naturalistic settings challenging. Although prior work has focused on improving model performance, it remains unclear whether wearable physiology supports a shared cross-individual mapping to subjective stress or whether this relationship is fundamentally person-specific.

We evaluated feature-based and deep-learning models across multiple physiological modalities using ecological momentary assessment (EMA) as the reference standard, comparing within- and between-individual modeling approaches. Within-individual models achieved modest but consistent improvements in stress detection, whereas between-individual models consistently failed to generalize, yielding negative *R*^2^ values despite multimodal fusion and high-capacity architectures.

Error analyses revealed regression to the mean, reduced sensitivity to high-stress states, and residual associations with general physiological activation, highlighting the limited stress specificity of wearable physiology. These findings suggest that wearable stress detection is fundamentally a personalized inference problem and that future systems should prioritize individual adaptation and contextual modeling over universal stress predictors.

## I. Introduction

Stress is a transdiagnostic process implicated in numerous psychiatric and somatic disorders[1], [2]. Reliable, unobtrusive detection using wearable devices could therefore enable scalable mental health monitoring. According to the transactional model of stress, stress is a subjective state arising from an individual’s appraisal of situational demands relative to available coping resources [3]. Because stress is not directly observable, detecting it from wearable physiology requires subjective self-reports as the reference standard, while physiology provides only indirect and noisy correlates of this latent experience [4]. Consequently, stress detection from physiology essentially a mapping problem between a latent experience and noisy biological signals [5], [6]. Because this mapping may vary substantially between individuals, models trained across participants often fail to generalize to unseen individuals, making reliable stress detection in everyday life an open challenge.

Laboratory measures such as cortisol assays, EEG, and standardized stress-induction protocols allow precise assessment under controlled conditions but are impractical for continuous real-world monitoring [7], [8]. Consequently, most field studies use ecological momentary assessment (EMA) as the reference standard for subjective stress [9]. However, repeated self-report is burdensome and is difficult to sustain over long periods, motivating the use of wearable physiology as a passive proxy.

Physiological signals measured by wearables primarily reflect general autonomic nervous system activity that is not necessarily specific to stress, making it difficult to distinguish stress from other high-arousal states such as physical activity or excitement [5], [6], [10], [11]. From an analytical perspective, while traditional approaches rely on features derived from research on stress physiology, it remains an open question whether modern machine learning methods can leverage raw sensor streams more effectively to uncover patterns that feature engineering may miss. Moreover, there is still no consensus on which physiological and motion signals, or their combinations, most reliably reflect subjective stress in daily life, as empirical findings across electrodermal, cardiovascular, thermal, and motion-based measures remain inconsistent [12], [13], [14]. Combining complementary physiological and motion modalities using machine learning could improve specificity by capturing multivariate patterns that are not evident in individual signals alone. Whether such multimodal representations can reliably generalize across individuals, however, remains an open question.

Prior work has largely focused on improving wearable-based stress detection through feature engineering, multi-modal sensing, and deep learning. While controlled or semi-controlled datasets such as WESAD provide labels linked to experimentally induced stress conditions [15], more recent studies have combined wearable sensing with EMA to study stress in daily life [16], [17], [18], [9]. However, it remains unclear whether multimodal wearable signals capture a common cross-individual mapping to subjective stress. Rather than treating poor between-individual performance solely as a modeling limitation, we investigate whether it reflects a more fundamental property of subjective stress—that its physiological correlates are inherently person-specific.

In this work, we provide a systematic analysis of wearable-based stress detection in naturalistic settings using a dataset of 894 participants with concurrent wearable-derived physiological measurements and EMA. We compare population-level and personalized models to assess generalization across individuals, evaluate feature-based and end-to-end learning approaches, and investigate the benefits of multimodal sensing. Additionally, we conduct a systematic analysis of model failure modes, examining when and why predictions break down across individuals, stress levels, and contexts. Specifically, we addressed the following research questions:

1. Can multimodal wearable physiological signals reliably detect real-life subjective stress states, as measured using ecological momentary assessment (EMA), within individuals?
2. If so, do the learned physiology–stress relationships generalize to unseen individuals, or are they fundamentally person-specific?
3. Do feature-based models provide more robust and generalizable representations of stress-related physiology than end-to-end models trained on raw signals?
4. Does combining multiple physiological modalities improve the detection of subjective stress relative to unimodal approaches?
5. Under what conditions do stress prediction models fail, and what factors (e.g., individual variability, stress distribution, and calibration behavior) explain these failures?

In this paper, we use the term “stress detection” rather than “stress prediction” to describe the inference of an individual’s current self-reported stress from concurrently acquired physiological signals. We avoid the term “prediction” because it is often interpreted as prospective forecasting of future stress states, which is beyond the scope of this work. Our objective is to examine whether physiological measurements can reliably infer an individual’s current subjective stress experience, rather than to forecast future stress or establish a causal relationship between physiological signals and subjective stress.

## II. Materials AND Methods

### A. Participants

Data were obtained from the Healthy Brain Study (HBS), a longitudinal cohort of 896 healthy adults aged 30-39 years conducted in Nijmegen, the Netherlands [19]. After preprocessing and quality control (Table I), 446 participants were included in the analyses. The study was approved by the Institutional Review Board of Radboud University Medical Center (2018-4894), and all participants provided written informed consent.

**TABLE I:** Participant inclusion and retention across study stages.

| Category | N |
| --- | --- |
| Initial cohort | 896 |
| *Assessment completion* |  |
| $\geq 1$ assessment period | 892 |
| $\geq 2$ assessment periods | 763 |
| All 3 assessment periods | 718 |
| *Final inclusion of participants* |  |
| who had wearable data | 883 |
| who had EMA data | 861 |
| who had matching wearable and EMA data (raw availability) | 851 |
| who had matching wearable and EMA data (after preprocessing) | 690 |
| who failed Quality Control by modality (participant fully removed) | 23 |
| who lacked enough EMA variation ($\geq 4$) | 241 |
| Total removed (accounting for overlap) | 244 |
| Final analytical sample | 446 |
| *EMA variability (final analytical sample)* |  |
| 4 unique stress labels | 173 (38.8%) |
| 5 unique stress labels | 178 (39.9%) |
| 6 unique stress labels | 79 (17.7%) |
| 7 unique stress labels | 16 (3.6%) |
| Mean EMA–physiology pairs per participant: 72.68 (SD = 39.03) |  |
| *Note.* Assessment completion was defined as the availability of either EMA or wearable data within a given measurement period. Physiological preprocessing and quality control were performed at the segment level, resulting in the removal of portions of data within participants rather than entire participants (see Section III). Participant-level exclusion ($N = 244$) was applied only after preprocessing, based on insufficient EMA variability and/or lack of valid EMA-physiology pairs for modeling. | |

For the present analyses, participants were required to have valid EMA-physiology pairs and sufficient within-person variability in stress ratings (*≥* 4 distinct levels) (see Table I).

### B. Study Design and Procedure

Data collection for the HBS took place between November 2020 and December 2023 [19]. For ecological monitoring, participants completed three six-day periods of EMA and wearable data collection, each separated by approximately three months. EMA was delivered via a smartphone application, with 10 prompts per day distributed at semi-random intervals within a predefined 12.5-hour daily sampling window (between 08:00 and 20:30), yielding up to 180 assessments per participant. During each monitoring period, participants continuously wore an Empatica E4 wristband (Empatica Inc., Milan, Italy [20]) on the non-dominant wrist, except during charging or water contact. Physiological data were collected via this wristband, a validated device for ambulatory stress research [21], [22], [23]. The device recorded electrodermal activity (EDA, 4 Hz), blood volume pulse (BVP, 64 Hz), 3-axis accelerometry (ACC, 32 Hz), and skin temperature (TEMP, 4 Hz). BVP and EDA were used as primary predictors of autonomic activity, while ACC and TEMP were included to quantify movement and peripheral thermoregulation. Physiological and EMA data were subsequently aligned using the 10-minute window preceding each EMA prompt to capture the physiological context of self-reported stress. Each EMA survey comprised 38 items assessing mood and context. Momentary mood was measured via 14 adjectives rated on a 7-point Likert scale (1 = not at all to 7 = very much). The single item ‘I feel stressed’ was used as a proxy for subjective stress. For secondary analyses, three additional items were included: “I feel cheerful,” “I feel relaxed,” and “I feel sad,” representing different regions of the valence-arousal spectrum. The unit of analysis was the EMA prompt (‘beep’).

### C. Data and Code Availability

All data are available from the Healthy Brain Study consortium upon request [https://www.healthybrainstudy.nl/en/data], subject to access restrictions. All preprocessing and analysis code is publicly available at [https://github.com/Selin99/Wearanize_Project] under an MIT license.

## III. Preprocessing

Physiological signals (Accelerometer (ACC), Temperature (TEMP), Blood Volume Pulse (BVP), and Electrodermal Activity (EDA)) were preprocessed with a dedicated pipeline to mitigate motion, contact, and environmental artifacts [24], [25] (See Figure 1 and Supplementary Material for full details).

**Fig. 1:**
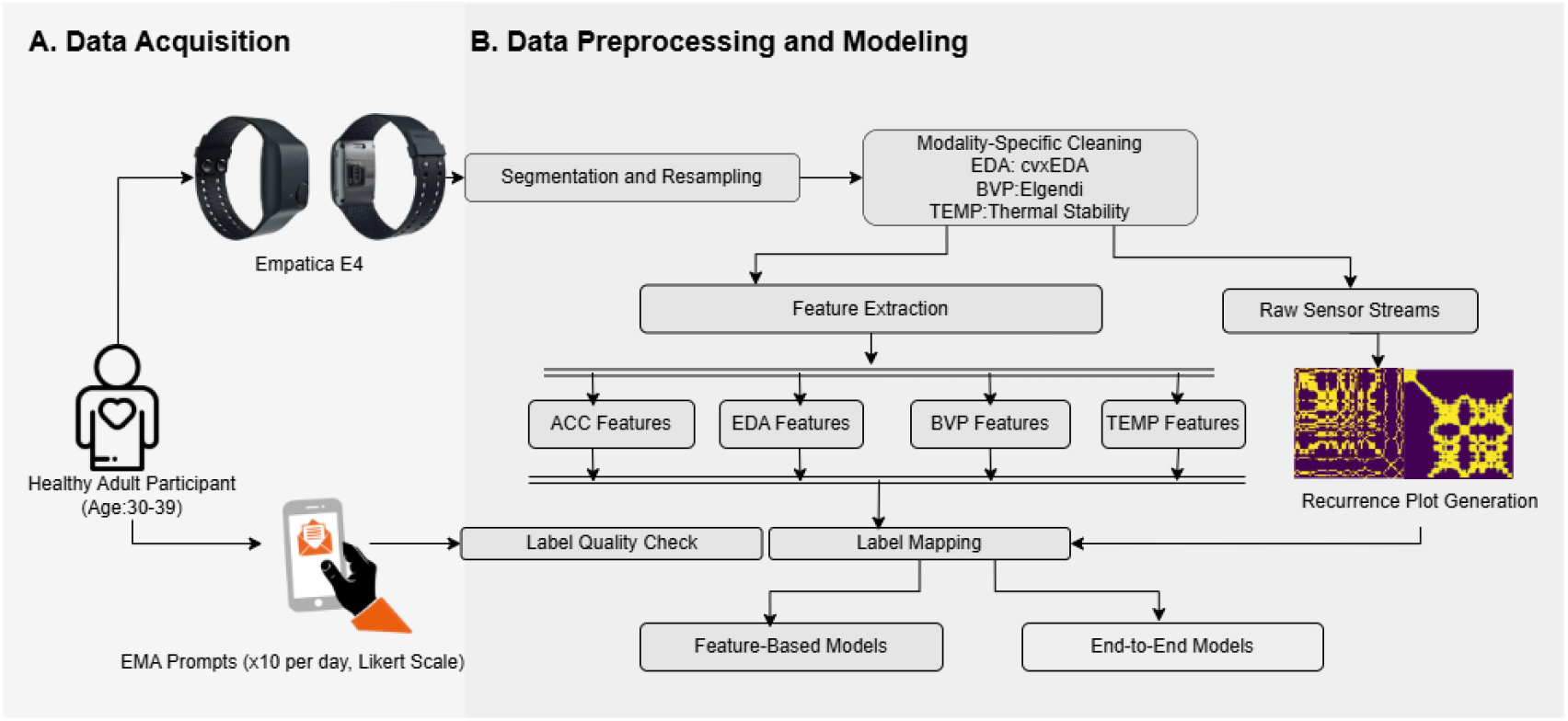
Overview of the preprocessing and feature extraction pipeline, including segmentation, quality control, and modeling representations.

Quality control removed noisy segments prior to feature extraction. Table II summarizes the extracted features.

**TABLE II:** Extracted physiological features grouped by sensing modality.

| Modality | Category | Features |
| --- | --- | --- |
| **ACC** | Time-domain | Mean, STD, RMS, MAD, IQR, skewness, kurtosis, energy, zero-crossing rate, jerk mean/STD. |
|  | Frequency-domain | Dominant frequency, spectral entropy, centroid, bandwidth, flatness, roll-off 85%, flux, variability. |
| **BVP/PPG** | Time-domain | MeanNN, SDNN, RMSSD, pNN50, BPM. |
|  | Frequency-domain | LF power, HF power, LF/HF ratio, total power. |
| | Nonlinear | Poincaré SD1/SD2, SampEn, ApEn, DFA $\alpha_1/\alpha_2$, Higuchi fractal dimension. |
| **TEMP** | Statistical | Mean, STD, AAD, min, max, IQR, MAD, skewness, kurtosis, proportion above mean. |
|  | Frequency-domain | Dominant frequency index, spectral entropy, centroid, bandwidth. |
| **EDA** | Time-domain | SCR peak count, mean peak amplitude, rise time, recovery time, mean SCL, AUC, amplitude variance, PPI mean/SD. |
|  | Frequency-domain | PSD LF, HF, LF/HF ratio, SCR skewness, kurtosis. |

Features were computed per segment and linked to corresponding EMA labels.

### a) End-to-end modeling

For end-to-end models, cleaned physiological signals were transformed into two-dimensional recurrence plots (RPs), which encode temporal dynamics as image representations suitable for convolutional neural networks (CNNs) [26]. RPs have been successfully applied to physiological time-series analysis, including ECG/PPG classification and wearable sensor data, by enabling CNNs to learn temporal patterns directly from recurrence structures [27], [28], [29], [30]. Furthermore, recurrence plots have been shown to provide an effective bridge between time-series analysis and image-based deep learning, enabling convolutional neural networks to learn temporal patterns directly from visual recurrence structures [29], [30]. Consequently, well-established convolutional neural network (CNN) architectures can be leveraged to automatically learn hierarchical spatial representations of temporal dynamics.

Recurrence plots were generated using the `pyts` implementation with embedding dimension *m* = 1, Euclidean distance, and a fixed recurrence rate of 30%. Signals were segmented into 2-minute windows, and segments with near-zero variance (SD *<* 0.001) were excluded. Additional implementation details are provided in the Supplementary Material.

Prior work has shown that deep CNNs operating on recur,rence plots achieve strong performance across a variety of physiological and time-series classification tasks [27], [28], [30].

### A. Statistical Analysis

To address the research questions stated in the Introduction Section I, analyses were organized around four objectives: evaluating cross-participant generalization, comparing feature-based and raw-signal representations, assessing multimodal integration, and characterizing model failure modes.

All analyses were performed in Python 3.11 using NumPy, pandas, scikit-learn, and TensorFlow. Feature-level analyses combined non-parametric statistics (Kruskal–Wallis H-test, Spearman’s rank correlation *ρ*) with dispersion and dependency metrics, including the Coefficient of Variation (CV) and Mutual Information (MI), to quantify variability and non-linear associations between physiological features and self-reported stress.

EMA stress was modeled as a continuous outcome to preserve ordinal information and avoid information loss associated with categorization. Furthermore, classification was not used as the primary formulation because binarizing or grouping stress ratings would discard information, introduce threshold dependence, and worsen class imbalance given the skew toward low-stress responses. Class imbalance in EMA stress ratings was addressed using class weighting during model training to account for skewed label distributions. For inter-user models, weights were computed based on the pooled distribution of EMA ratings across all training participants. For intra-user models, weights were computed separately within each participant to reflect individual stress distributions.

#### a) Model architectures

Motivated by the heterogeneity in modeling approaches and performance reported in prior work [31], we systematically evaluate whether detection of subjective stress states from physiological data constitutes a learnable cross-individual mapping under realistic conditions. The architectures included linear, ensemble, and deep learning paradigms to ensure that conclusions are not driven by a single modeling framework. Model families were selected to test whether increasing representational capacity can compensate for the absence of shared physiological-stress structure.

For feature-extracted data, we trained Random Forests, Gradient-Boosted Trees (XGBoost), Multi-Layer Perceptrons (MLP), one-dimensional Convolutional and Recurrent Neural Networks (1D-CNN, LSTM, RNN), and linear baselines (Ridge, LASSO). For end-to-end models, we evaluated 2-D deep convolutional architectures including AlexNet, ResNet, DenseNet, and GoogLeNet.

These architectures have been widely applied to recurrence-plot representations of physiological and time-series data, including ECG/PPG classification and wearable sensor analysis, where they have demonstrated strong performance in learning discriminative temporal patterns directly from image-based representations [27], [28], [29], [30]. Evaluating architectures with progressively greater representational capacity allowed us to assess whether limitations in stress prediction arise from insufficient model complexity or from the absence of a shared physiological-stress mapping across individuals.

Hyperparameters were optimized using nested cross-validation (grid-search). All evaluation and statistical analyses were performed at the participant level to account for correlated observations and prevent data leakage. Cross-validation splits were therefore defined by participant, while intra-user analyses used non-overlapping temporal segments. Physiological windows corresponding to the same EMA prompt (“beep”) were assigned to the same split to ensure complete separation between training and test data.

#### b) Modeling Strategy

Three complementary modeling strategies were evaluated:

- **Inter-user feature models (LOSO):** Feature-based models were trained on all participants except one and evaluated on the held-out participant using leave-one-subject-out (LOSO) cross-validation. Performance was averaged across folds to assess cross-individual generalization.
- **End-to-end models (LOSO):** Deep learning models were trained directly on recurrence-plot representations of physiological signals using the same LOSO procedure, enabling comparison between feature-based and end-to-end representations.
- **Intra-user feature models:** Personalized feature-based models were trained and evaluated on non-overlapping, temporally ordered train/test splits (70%/30%) within each participant to assess individual-specific physiology-stress relationships.

Importantly, the LOSO and intra-user approaches address different generalization targets. LOSO models evaluate unseen participants, whereas intra-user models evaluate unseen observations from the same participant. While personalized models benefit from participant-specific physiology, LOSO models leverage substantially more training data. Consequently, the comparison reflects a trade-off between data quantity and personalization, making the two approaches complementary rather than directly comparable. Intra-user analyses were conducted only for feature-based models; raw-signal architectures were excluded from within-participant modeling due to insufficient data length and variability after preprocessing and exclusion criteria.

#### c) Performance metrics

Model performance was quantified using the coefficient of determination (*R*^2^), mean absolute error (MAE), and root mean squared error (RMSE). Ninety-five percent confidence intervals (CI) were estimated via nonparametric bootstrap resampling across folds.

#### d) Statistical testing

To evaluate whether model performance exceeded chance, we conducted nonparametric permutation tests, shuffling EMA labels, taking into account the participant and temporal dynamics, to generate null distributions for *R*^2^, MAE, and RMSE. Pairwise model comparisons were performed with paired *t*-tests; Wilcoxon signed-rank tests were additionally used for robustness to non-normality. All *p*-values were corrected for multiple comparisons using the Benjamini-Hochberg false discovery rate (FDR). Effect sizes were quantified with Cohen’s *d*, computed as the mean difference divided by the pooled standard deviation (See Supplemental Material for Details).

To characterize model failure modes, we analyzed errors at multiple levels: high-stress detection, calibration bias (systematic over or underestimation of stress levels), associations between performance and data availability, construct specificity relative to other affective EMA items, residual-feature correlations, and the effect of temporal aggregation from momentary to daily and weekly predictions.

#### 1) Baseline Performance and Label Distribution

Model performance was benchmarked against constant predictors based on the mean training-set stress score (global mean for population models; participant-specific mean for personalized models). Improvements over these baselines indicate added value beyond mean-level prediction. For *R*^2^, zero corresponds to the mean predictor, and negative values indicate worse performance than mean-level prediction within the evaluation data.

At the participant level (see Figure 2 and Table III), predicting the midpoint yielded a mean MAE of 1.64 (*±*0.50) with substantial interindividual variability (0.88–2.41). Using each participant’s median EMA score as a constant predictor markedly reduced error to 1.05 (*±*0.26), establishing a strong within-person reference. Hence, intra-user models were required to outperform this ~1.05 MAE baseline to demonstrate added predictive value.

**Fig. 2:**
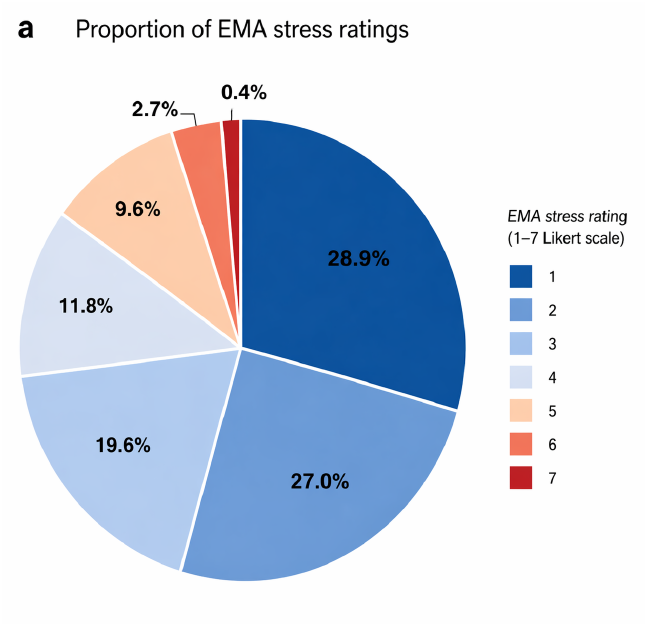
Proportion of EMA stress ratings (1-7 Likert scale) across all valid participants (*N* = 446). The higher empirical baseline reflects the strong left-skew of EMA ratings toward low stress values (1-3).

**TABLE III:**
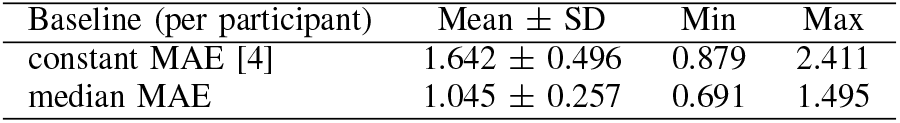
Participant-level constant baselines (*n* = 446). The constant midpoint baseline always predicts 4 (on Likert Scale), whereas the median baseline predicts each participant’s median EMA stress rating.

| Baseline (per participant) | Mean ± SD | Min | Max |
| --- | --- | --- | --- |
| constant MAE [4] | $1.642 \pm 0.496$ | 0.879 | 2.411 |
| median MAE | $1.045 \pm 0.257$ | 0.691 | 1.495 |

## IV Results

Results are organized around the four research questions: cross-participant generalization, feature-based versus raw-signal modeling, multimodal integration, and model failure modes.

### A. Physiological Variability and Stress Coupling

Because model performance is bounded by the strength and consistency of physiological-stress coupling, we first quantified signal variability and feature relevance at both intra-user (within-individual) and inter-user (across-individual) levels.

Within individuals, physiological features exhibited substantial heterogeneity in both variability and stress coupling (Fig. 3). Feature-level dispersion ranged from 0.8% to 63% (median = 14.2%), with temperature and heart rate showing relatively low variability (median CV *≈* 5–12%) and PRV and EDA features substantially higher variability, frequently exceeding 50% and, for several features (e.g., LF/HF and EDA amplitude metrics), 100%. Despite this variability, physiological features exhibited measurable within-subject associations with stress. Mutual information ranged from *≈* 0.30 to 1.25 across features (e.g., RMSSD *≈* 0.84, SDNN *≈* 0.85, mean SCL *≈* 1.25, SCR amplitude *≈* 1.22), while more than 90% of participants showed significant stress-related differences for key PRV and EDA features (RMSSD 94.2%, SDNN 95.2%, mean SCL 96.2%), with moderate effect sizes (median *η*^2^ *≈* 0.07–0.15).

**Fig. 3:**
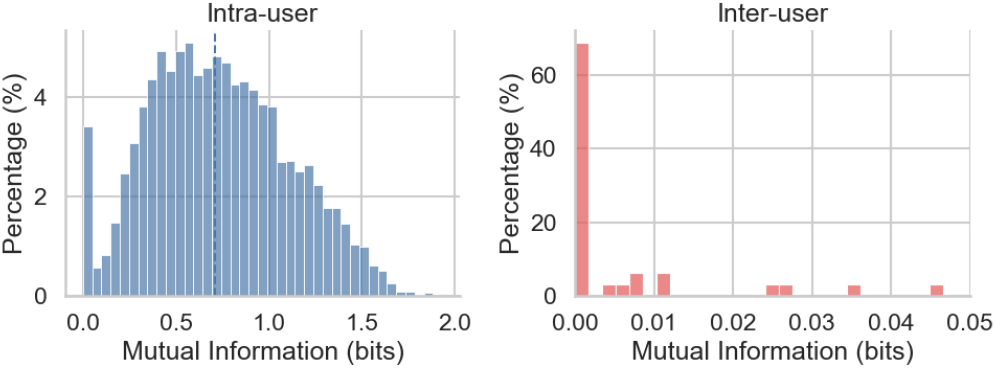
Mutual information (MI) was computed between each physiological feature and EMA stress ratings. For intra-user analyses, MI was estimated separately within each participant and then aggregated across participants. For inter-user analyses, MI was estimated using pooled observations across participants. The dashed vertical line indicates the median mutual information across all feature-participant combinations. Mutual information (MI) distributions show that physiological features exhibit meaningful dependency with stress within individuals, whereas inter-user MI values collapse toward zero.

In contrast, physiology–stress coupling was effectively absent at the population level. Mutual information was near zero for all features (median MI *≈* 0.00–0.02), Kruskal–Wallis tests were non-significant (*p >* 0.3), and effect sizes were negligible (median *η*^2^ *<* 0.005; maximum *η*^2^ *≈* 0.006). Notably, the most informative within-subject features (PRV and EDA; MI *>* 0.8, *η*^2^ *>* 0.1) also exhibited high variability, suggesting that informative physiological signals are inherently dynamic but poorly standardized across individuals. However, variability alone was insufficient, as several high-variance accelerometer features showed little stress-related information. Overall, these findings indicate that informative physiological patterns are largely individualized and collapse at the population level, motivating personalized modeling in subsequent analyses.

### B. Inter-user Generalization: Feature-based vs Raw Models

To evaluate whether stress can be predicted at the population level, we evaluated feature-based models and end-to-end deep learning models under a leave-one-subject-out (LOSO) framework and tested whether representation learning can overcome the lack of shared physiological structure (See Supplementary Material Section 3 and Section 4 for full model training details and statistical testing results).

Across all models and input representations, inter-user performance failed to explain meaningful variance in stress, with uniformly negative *R*^2^ values (Table IV), indicating performance below a mean-level predictor. Feature-based models achieved relatively consistent errors (MAE *≈* 1.16–1.22; RMSE *≈* 1.36–1.46), with EDA+TEMP performing best (*R*^2^ *≈ −*0.30). Although all models improved upon the constant baseline (MAE *≈* 1.64; Section III), none explained variance beyond mean-level prediction. Multimodal fusion yielded only marginal improvements, and ensemble methods consistently outperformed linear and deep sequence models (Cohen’s *d* = 0.3–0.5). End-to-end CNNs performed worse overall (MAE *≈* 1.17–1.70; RMSE *≈* 1.35–1.86; *R*^2^ *≈ −*0.24 to *−*0.78). Although ResNet generally outperformed other CNNs and DenseNet performed worst, architectural differences did not alter the overall pattern, and multimodal raw-signal models likewise failed to improve generalization.

**TABLE IV:**
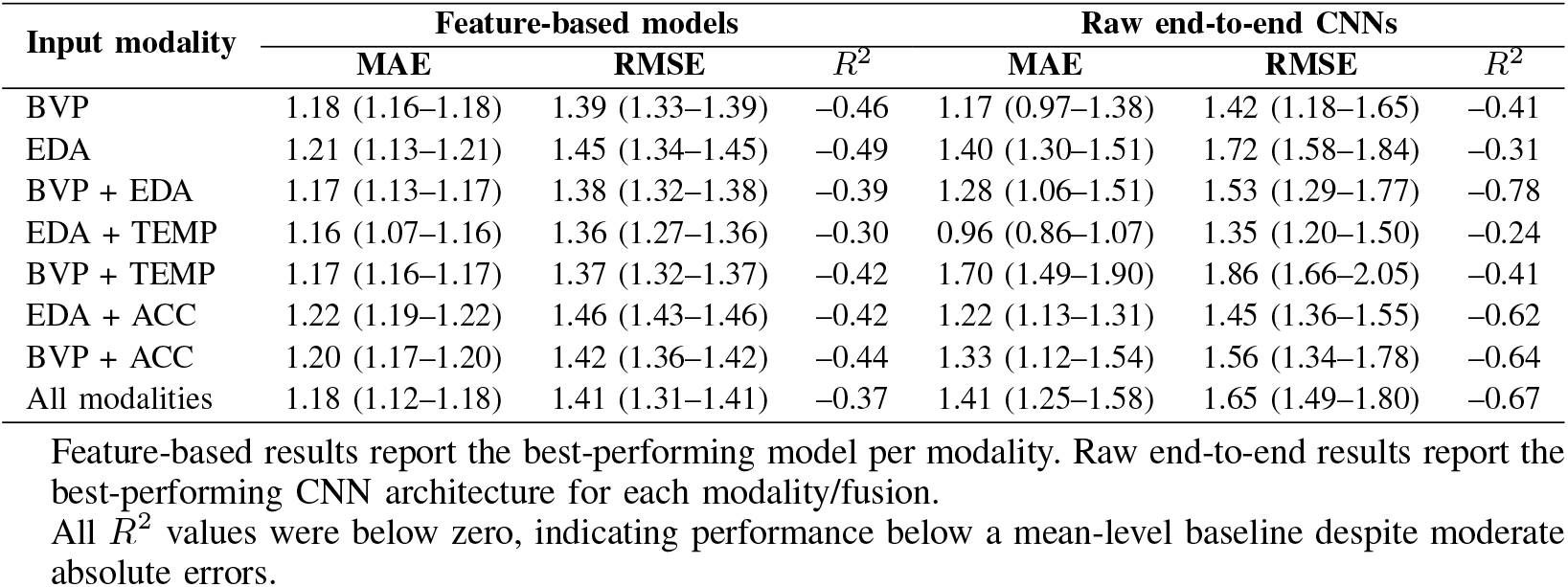
Inter-user LOSO performance for feature-based and raw end-to-end models predicting EMA stress ratings. Values are mean with 95% CI.

**TABLE V:**
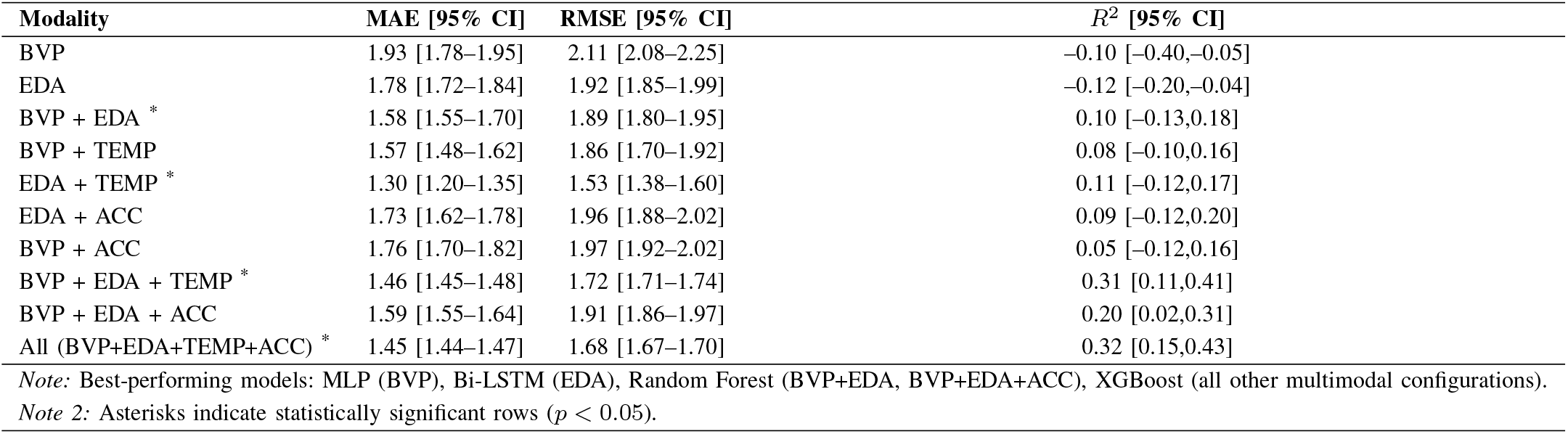
Summary of intra-user results with feature-extracted models (Target = EMA). Values correspond to the best-performing model per modality (selected by mean *R*^2^).

Overall, feature-based models consistently outperformed end-to-end approaches, suggesting that engineered features improve signal-to-noise ratio. However, uniformly negative *R*^2^ values and non-significant permutation tests (*p >* 0.05) indicate that neither representation learned a shared physiology-stress mapping across individuals. The combination of moderate MAE and negative *R*^2^ further suggests regression toward the population mean rather than accurate tracking of individual stress fluctuations.

### C. Feature Extracted Intra-user Models

We next evaluated intra-user models trained and tested within individuals using the same feature representations as in the inter-user experiments (See Supplementary Material Section 5 for full model training details and statistical testing results).

Intra-user models trained on feature-extracted summaries showed substantially improved performance compared to interuser models. Unimodal models based on BVP and EDA showed limited predictive value, with negative *R*^2^ values (*≈ −*0.10 to *−*0.12), indicating no improvement over a mean-level baseline, eventhough we observed performance gains over the inter-user alternatives. Despite moderate absolute errors and significant differences between model classes (Wilcoxon *p <* 0.01, *d >* 0.8), these models failed to capture within-individual variation in stress. Bimodal models yielded modest improvements, with small positive explained variance (*R*^2^ *≈* 0.05–0.11). Among these, EDA+TEMP consistently performed best, suggesting that thermoregulatory signals provide complementary information to electrodermal

activity. However, gains remained limited, indicating that two modalities are insufficient to robustly capture stress dynamics. Across modalities, tree ensembles (especially XGBoost) consistently outperformed linear and deep sequence models (all *d >* 1.0). Substantial improvements emerged when combining three modalities. In particular, BVP+EDA+TEMP achieved *R*^2^ *≈* 0.31 with reduced error, representing a marked increase over bimodal models. The full multimodal configuration (BVP+EDA+TEMP+ACC) further improved performance to *R*^2^ = 0.32, although gains beyond tri-modal fusion were modest. Although intra-user models achieved positive *R*^2^, their MAE remained above the participant-median constant baseline (median MAE *≈* 1.05; Section III), indicating that they captured variance but did not outperform a strong within-person absolute-error baseline.

Within-subject calibration allows models to learn individual-specific physiology-stress relationships, including baseline differences and reactivity patterns that are not captured by population-level models. Without such calibration, physiological signals show limited generalizability across individuals. Overall, these findings reveal a consistent pattern across analyses. Feature-level results showed that physiological signals contain meaningful information about stress within individuals, as also reflected by moderate mutual information and significant within-subject effects. However, this structure largely disappears at the population level. On the other hand, even within individuals, predictive performance remained limited, with substantial unexplained variance and strong heterogeneity across participants when the failure points analysed (See Section IV-D2).

### D. Intra-User Models: Feature Importance, Participant Heterogeneity, Failure Modes, Construct Specificity, and Temporal Aggregation

Subsequent analyses focus on the best-performing intra-user models to characterize feature relevance and sources of prediction error.

#### 1) Feature Importances

To quantify the contribution of individual physiological features to stress prediction, feature importance was estimated using SHapley Additive exPlanations (SHAP) values derived from the best-performing intra-user XGBoost models. Feature importance was computed as the mean absolute SHAP value and subsequently averaged across participants. Feature importance analysis indicated that electrodermal activity (EDA) and peripheral temperature features contributed most strongly to stress prediction (Fig. 4). In particular, tonic skin conductance level (SCL) and phasic SCR dynamics were consistently ranked among the top predictors, followed by temperature-derived features. These findings are consistent with established stress physiology, where sympathetic activation modulates sudomotor activity and peripheral vasoconstriction [32], [33], [34].

**Fig. 4:**
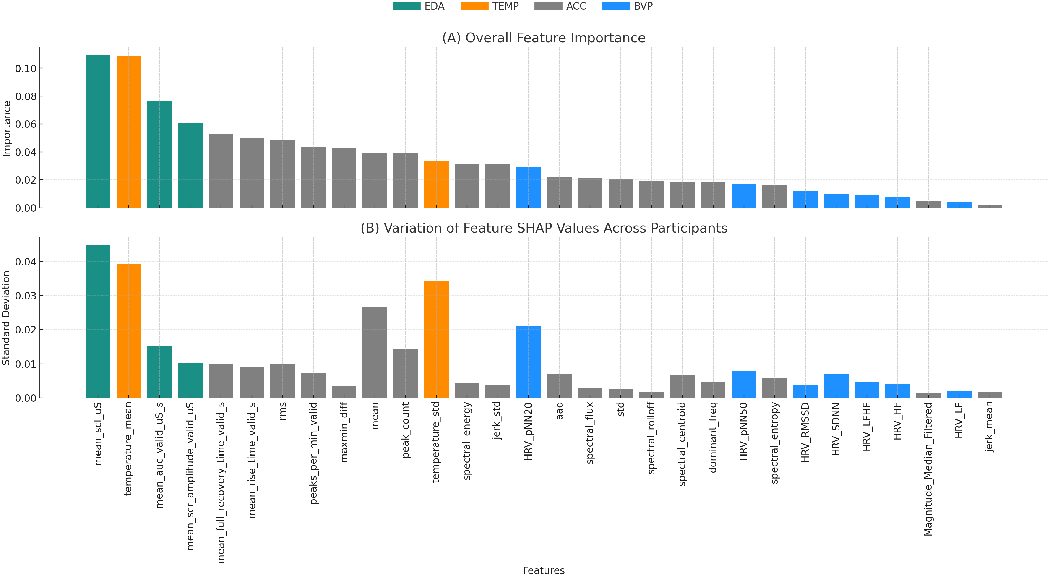
Feature importance and inter-individual variability derived from SHAP analyses of the best-performing intra-user XGBoost models. (A) Mean absolute SHAP values averaged across participants, indicating overall feature importance. (B) Standard deviation of SHAP values across participants, indicating inter-individual variability in feature importance.

In contrast, PRV-derived features contributed less to model performance. This may reflect measurement limitations, as cardiovascular responses to stress are more indirect and wearable PPG signals are very susceptible to motion artifacts and noise [35], and the loss of PRV information resulting from the exclusion of low-quality PPG segments during preprocessing.

Notably, the features with the highest predictive importance also exhibited the greatest inter-individual variability, suggesting that the most informative physiological signals are also the least standardized across participants. Features such as temperature variability exhibited relatively low average importance but high between-participant variability in SHAP values, suggesting that they are informative for a subset of individuals while contributing little to no prediction in most participants. This pattern is consistent with substantial inter-individual heterogeneity in physiological stress responses. From a physiological perspective, stress-induced sympathetic activation can trigger peripheral vasoconstriction, leading to reductions in skin temperature and altered thermal dynamics. However, the strength and direction of these responses vary considerably across individuals due to differences in autonomic reactivity, thermoregulatory regulation, vascular responsiveness, environmental influences, and behavioral factors. Furthermore, temperature fluctuations are not specific to stress and can also arise from physical activity, ambient temperature changes, circadian processes, emotional arousal, and other forms of autonomic activation. As a result, temperature-derived features may serve as strong stress indicators for some participants but fail to generalize across the broader population.

#### 2) Participant Heterogeneit

Prediction performance varied substantially across individuals (Fig. 5), with only subset of users achieving both low error and meaningful explanatory power.

**Fig. 5:**
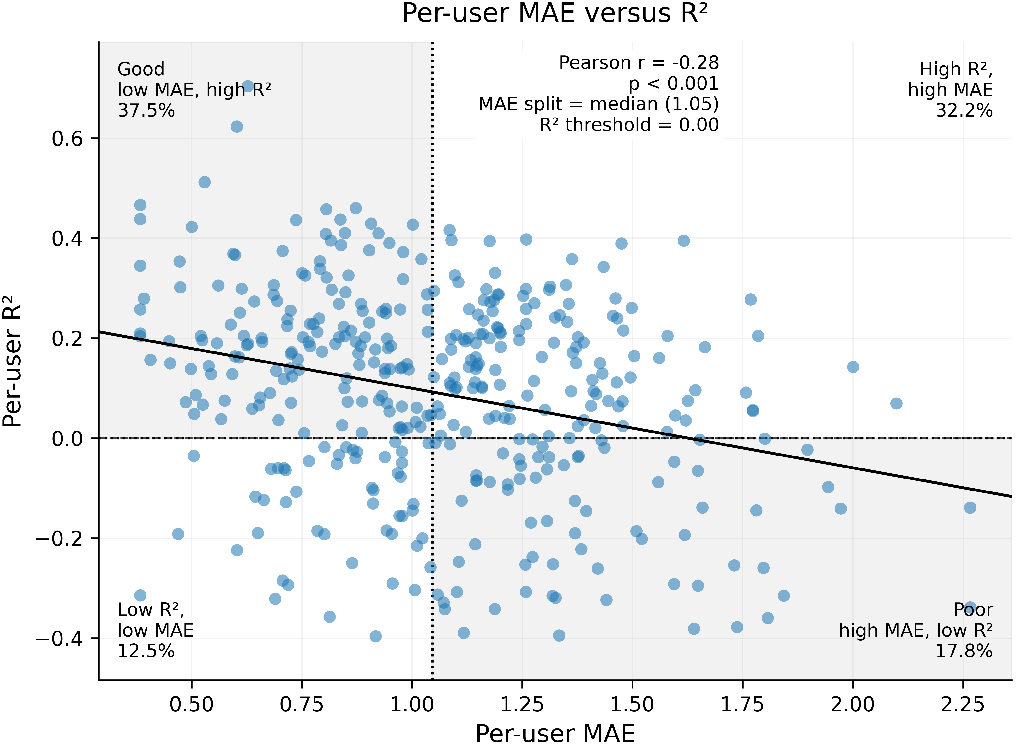
Relationship between per-participant prediction error (MAE) and explanatory power (*R*^2^). Each point represents an individual user. The vertical dashed line indicates the median MAE for intra-user analysis (see Section 2 for details), and the horizontal dashed line indicates *R*^2^ = 0, corresponding to performance equal to a naïve baseline. Shaded regions highlight users with relatively favorable (low MAE, high *R*^2^) and unfavorable (high MAE, low *R*^2^) performance.The modest association between MAE and *R*^2^ (*r* = 0.29) highlights that low prediction error does not necessarily imply successful tracking of stress dynamics. Many participants exhibited relatively low MAE despite near-zero explained variance, indicating that models frequently approximated individual baseline stress levels while failing to capture moment-to-moment fluctuations.

Model performance varied substantially across individuals (mean *R*^2^ = 0.3, SD = 0.20), with *≈* 30% of users exhibiting negative *R*^2^ values. Only 31% of users achieved *R*^2^ *>* 0.3, indicating that strong predictive performance was limited to a subset of participants. This distribution highlights that the average model performance was driven by a subset of participants with stronger physiological-stress coupling. Together, these findings indicate that stress responses vary both in magnitude and in which physiological systems dominate per person, making personalized modeling a necessity rather than a design choice.

##### a) Characterizing Prediction Failure and Error Structure

We investigate why the model performs poorly for certain individuals, focusing on high-stress misclassification, predictability, contextual modulation, and residual structure. The following analysis were performed on the best-performing within-individual models.

##### b) High-Stress Classification Performance

High stress was defined using a user-specific upper quartile threshold (*q*_*high*_ = 0.75), corresponding to the top 25% of reported stress values within each individual, relative to each individual’s baseline rather than using a fixed absolute cutoff. Sensitivity analyses using alternative within-person thresholds (e.g., top 20% and 30%) yielded qualitatively similar results, indicating that conclusions are not driven by the specific choice of cutoff. Predictions were binarized into high vs. non-high stress, and confusion matrices were computed per user. From these, recall, precision, false positive rate (FPR), and F1 scores were derived and summarized across users (Table VI).

**TABLE VI:** Per-user high-stress detection performance.

| Metric | Mean | Median | SD | IQR |
| --- | --- | --- | --- | --- |
| Recall | 0.55 | 0.55 | 0.12 | 0.50–0.63 |
| False Positive Rate | 0.46 | 0.50 | 0.26 | 0.33–0.57 |
| Precision | 0.76 | 0.75 | 0.21 | 0.63–0.87 |
| F1 Score | 0.64 | 0.67 | 0.12 | 0.57–0.71 |

Across participants, recall was moderate (mean = 0.55), while precision was higher (mean = 0.76), indicating that predicted high-stress events were often correct but frequently missed. The false positive rate remained substantial (mean = 0.46), reflecting variability across users. This asymmetry (*precision > recall*) indicates that the model is conservative, tending to under-detect high-stress episodes. As a result, false negatives dominate, suggesting reduced sensitivity to extreme stress states.

To determine whether this tendency reflects a systematic prediction bias, we next examined model calibration. In this context, calibration refers to the degree to which predicted stress levels correspond to reported stress levels across the full range of the outcome. Poor calibration, particularly compression of predictions toward the mean, would be expected to systematically underestimate high-stress observations and could therefore explain the observed excess of false negatives.

Calibration analysis further demonstrated strong compression of predicted values toward the mean. The median calibration slope was low (*β*_1_ *≈* 0.2), substantially underestimating the magnitude of fluctuations. The model captures relative variation in stress but, it has limited ability to estimate absolute stress levels.

Consistent with this pattern, prediction error increased markedly with stress intensity (Fig. 6A), rising from approximately 1.0 in low-to-moderate stress ranges to 2.75 in the highest quantile. This suggests that while the model captures relative changes in stress (as reflected by moderate *R*^2^), it fails to accurately represent the magnitude of these changes. Despite the use of class weighting during training, the model exhibited pronounced regression-to-the-mean behavior, indicating that this effect cannot be fully explained by class imbalance alone. Instead, these results suggest that the underlying physiological signal associated with higher stress levels (values outside the homeostatic range) is weaker, more variable, or less consistently captured by the available features. This limits the model’s ability to accurately represent extreme stress states, leading to systematic underestimation.

**Fig. 6:**
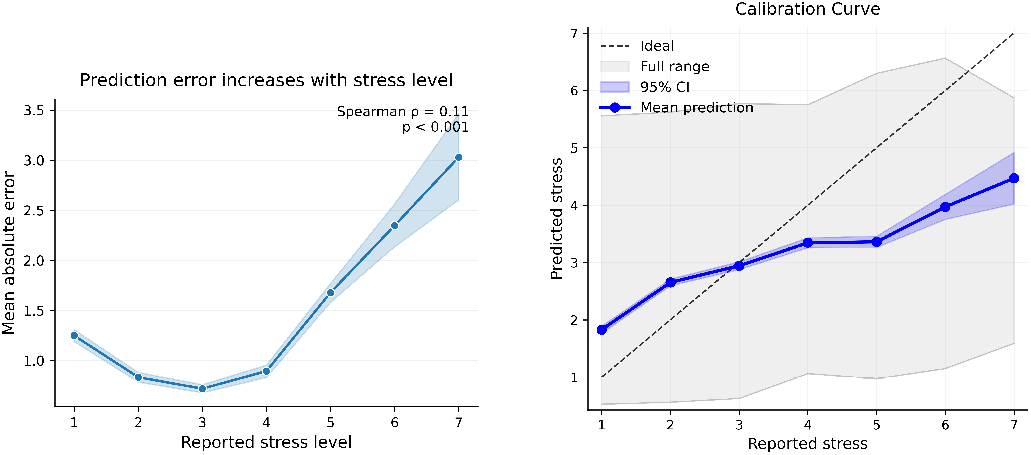
Prediction error and calibration of the stress model. (A) Mean absolute error increases with reported stress level, indicating reduced accuracy at higher stress levels. (B) Calibration curve showing systematic compression of predictions toward the mean, with overestimation at low stress and underestimation at high stress.

##### c) Role of data quantity and class imbalance in model performance

To assess the potential influence of label distribution on model performance, class imbalance was quantified separately for each participant as the proportion of observations belonging to the high-stress class relative to the total number of observations. Spearman correlations were then computed between participant-level (*R*^2^) values and (i) the total number of EMA observations, (ii) the participant-specific class imbalance measure, and (iii) the absolute number of high-stress observations (*n*_1_).

Neither total data quantity nor class imbalance showed a meaningful association with performance. The correlation between *R*^2^ and the number of observations was weak and non-significant (*ρ* = 0.11, *p* = *−*0.066), and class imbalance exhibited only a weak negative trend (*ρ* = −0.09, *p* = 0.147). These results indicate that neither increasing the overall amount of data nor altering the relative class distribution substantially improves predictive accuracy.

In contrast, the number of high-stress observations (*n*_1_) showed a strong positive association with model performance (*ρ* = 0.69, *p <* 0.001). This suggests that performance is primarily driven by exposure to informative events rather than total data volume.

These findings are consistent with prior work by Smets et al. [17], which reported that high-performing individuals tended to have more higher reported stress levels and slightly higher response rates. In line with this, we also observe only a weak association between total data quantity and performance.

Notably, the absence of a strong imbalance effect may indicate that the class-weighting strategy applied during model training partially mitigated the impact of unequal label distributions. However, because weighted and unweighted models were not directly compared, this interpretation should be considered tentative. Smets et al. further reported that high-performing individuals tended to have greater exposure to stress-related events. Our results extend this observation by showing that the number of high-stress observations is strongly associated with performance at the individual level, suggesting that learning is driven by exposure to informative physiological states rather than overall data volume.

Importantly, these findings should be interpreted in light of the limited observation window. Data were collected over a three-week period and split into training and testing sets (70:30), which constrains the total amount of data available for model learning. It is therefore possible that longer monitoring periods could improve performance by providing more stable estimates of individual physiological patterns and greater exposure to stress events.

##### d) Predicted Stress and Competing Affective Constructs

The primary objective of this analysis is to assess the construct validity of the predicted stress signal, specifically whether model outputs align uniquely with subjective stress or instead reflect broader affective dimensions such as valence or general arousal. Residuals were defined as the difference between predicted and reported stress (residual = *y*_pred_ *− y*_true_), such that a positive Spearman correlation (*ρ >* 0) indicates systematic overestimation of stress as the feature increases, whereas a negative correlation (*ρ <* 0) indicates systematic underestimation.

Residual-feature analysis (see Table VII) revealed that prediction errors exhibited systematic dependencies on affective variables, indicating non-random structure. Positive valence, positive arousal (cheerful) showed a substantial positive association with residuals (mean *ρ* = 0.42, median *ρ* = 0.45, SD = 0.28; 29.7% significant), indicating systematic overestimation of stress during positive arousal states. This finding is consistent with previous observations that wearable physiological signals alone often fail to distinguish stress from other high-arousal emotional states, particularly those with positive valence [9]. Similarly, relaxed states exhibited a positive association with the residuals (mean *ρ* = 0.29, median *ρ* = 0.33, SD = 0.31; 19.0% significant), and negative valence (sad) showed a moderate negative association with the residuals (mean *ρ* = *−*0.26, median *ρ* = *−*0.31, SD = 0.32; 12.9% significant), suggesting that the model partially captures but does not fully align with affective dimensions.

**TABLE VII:** Per-participant residual-feature correlations. Signed residual correlations indicate the direction of bias (*predicted − reported* stress), whereas absolute residual correlations indicate associations with error magnitude. Fraction significant denotes the proportion of users with nominally significant per-user Spearman correlations (*p <* 0.05).

| Variable | Mean $\rho_{\text{signed}}$ | Median $\rho_{\text{signed}}$ | SD $\rho_{\text{signed}}$ | Mean $\rho_{\text{abs}}$ | Median $\rho_{\text{abs}}$ | SD $\rho_{\text{abs}}$ | Frac. sig. signed | Frac. sig. abs |
| --- | --- | --- | --- | --- | --- | --- | --- | --- |
| Cheerful | 0.4153 | 0.4542 | 0.2811 | -0.0927 | -0.0839 | 0.2998 | 0.2966 | 0.0646 |
| Relaxed | 0.2886 | 0.3293 | 0.3110 | -0.0721 | -0.0861 | 0.2779 | 0.1901 | 0.0456 |
| Sad | -0.2623 | -0.3057 | 0.3190 | 0.0109 | 0.0000 | 0.3071 | 0.1293 | 0.0532 |
| temperature std | 0.0168 | 0.0070 | 0.2788 | 0.0580 | 0.0373 | 0.2936 | 0.0304 | 0.0380 |
| jerk std | -0.0069 | -0.0209 | 0.3150 | 0.0569 | 0.1062 | 0.2977 | 0.0684 | 0.0570 |
| spectral energy | -0.0246 | -0.0195 | 0.3117 | 0.0442 | 0.0858 | 0.2979 | 0.0722 | 0.0532 |
| peak count | -0.0156 | 0.0228 | 0.2874 | 0.0437 | 0.0617 | 0.2918 | 0.0456 | 0.0494 |
| rms | -0.0077 | 0.0067 | 0.3035 | 0.0408 | 0.0364 | 0.2999 | 0.0456 | 0.0342 |
| mean temperature | 0.0079 | 0.0295 | 0.2976 | -0.0199 | -0.0350 | 0.3014 | 0.0570 | 0.0418 |
| dominant frequency | 0.0021 | 0.0097 | 0.2885 | 0.0176 | 0.0437 | 0.2833 | 0.0418 | 0.0266 |
| jerk mean | -0.0233 | 0.0050 | 0.3117 | -0.0054 | 0.0205 | 0.2832 | 0.0684 | 0.0380 |
| spectral entropy | -0.0055 | 0.0059 | 0.2804 | 0.0016 | 0.0000 | 0.2858 | 0.0456 | 0.0418 |

In contrast, motion and signal-related features exhibited near-zero signed correlations, including jerk variability (jerk std: mean *ρ* = *−*0.01), spectral energy (mean *ρ* = *−*0.02), peak count (mean *ρ* = *−*0.02), and RMS (mean *ρ* = *−*0.01), with low fractions of significant associations (7%). Temperature-related variables also showed negligible effects (temperature std: mean *ρ* = 0.02, temperature mean: mean *ρ* = 0.01). These findings indicate that prediction errors are not primarily driven by motion artifacts or environmental confounds.

##### e) Temporal Aggregation: From Momentary to Long-Term Stress Tracking

We assess whether predicted and reported stress co-vary within individuals across three temporal scales: beep (momentary), day, and week. Predictions were not refit at coarser time scales. Instead, beep-level predicted and reported stress values were averaged within daily and weekly bins.

Temporal aggregation had mixed effects on predictive accuracy. At the population level, MAE did not improve with aggregation ([1.13 at the beep level to 1.11 at the day level and 1.10 at the week level]. In contrast, RMSE decreased, from 1.41 at the beep level to 1.18 at the day level and 1.29 at the week level, indicating that aggregation reduced high-frequency variability but did not improve absolute prediction accuracy. At the individual level, the effect of aggregation was heterogeneous, as shown in Figure 7. Specifically, 12.9% of users experienced increased error (i.e., negative gain), with some cases showing substantial deterioration (minimum gain −2.78), indicating that aggregation at this timescale does not consistently improve performance. In contrast, week-level aggregation was more stable, yielding a larger mean MAE reduction of 0.53 (median 0.48), and only 1% of users showed worsening performance (minimum gain −0.23). Together, these findings suggest that longer-term averaging smooths noise more effectively than day-level averaging, but does not fundamentally resolve model error. This pattern suggests that the model benefits from temporal smoothing, particularly at the week level, but remains limited by systematic bias rather than purely short-term noise.

**Fig. 7:**
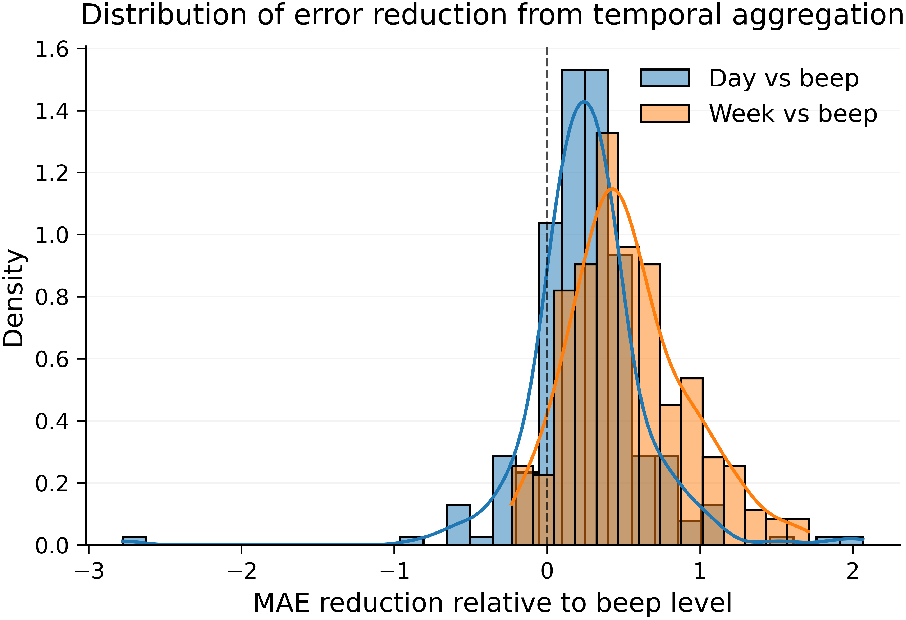
The figure shows the distribution of per-participant changes in mean absolute error (MAE) when aggregating predictions from the beep level to day and week levels. Positive values indicate improved performance (lower MAE), while negative values indicate worsening performance relative to the beep level. Week-level aggregation resulted in larger and more consistent improvements (*mean*Δ*MAE* = 0.53, *median* = 0.48) compared to day-level aggregation (*mean*Δ*MAE* = 0.26, *median* = 0.25). The vertical dashed line at zero denotes no change in error. Distributions are shown as normalized densities to facilitate comparison of their shapes.

**Fig. 8:**
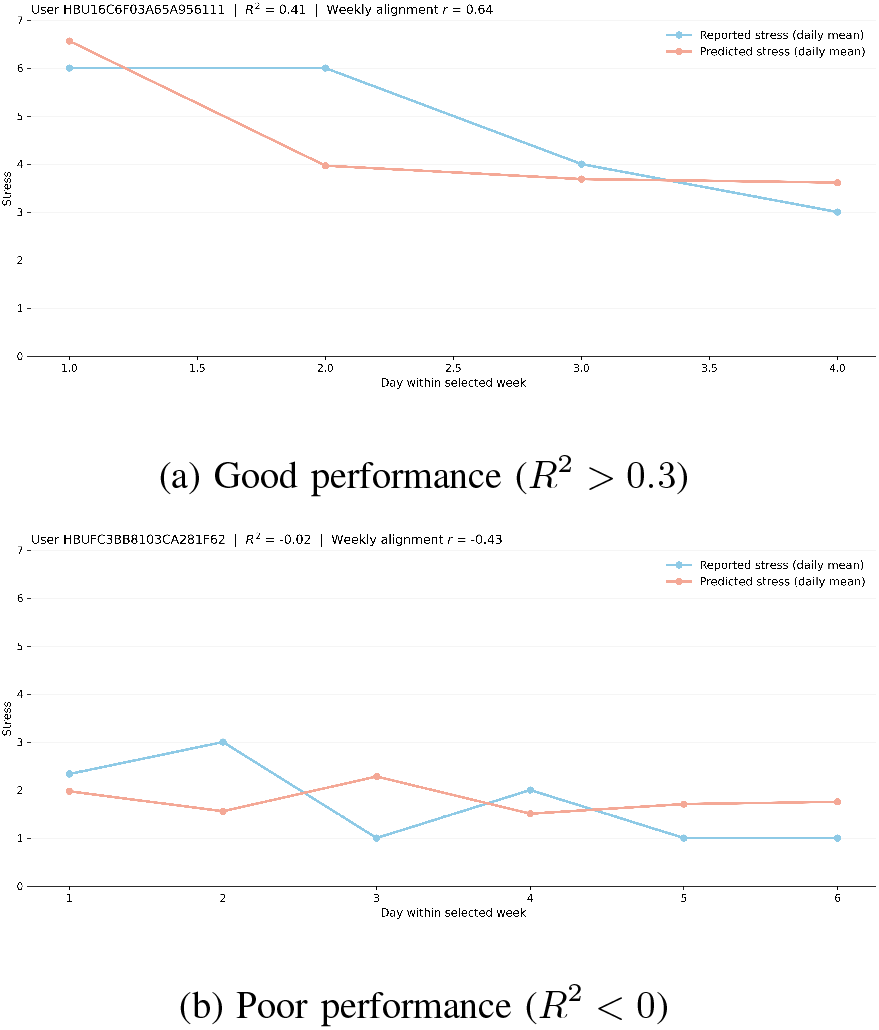
Representative examples of model performance across users. Each panel shows daily averaged reported and predicted stress within a selected week. Titles report per-participant predictive performance (*R*^2^) computed across the full dataset and the within-week correlation (*r*).

Temporal aggregation improved performance metrics, however, aggregation also makes the prediction problem substantially easier by removing much of the moment-to-moment variation that the models are intended to capture. Consequently, improved week-level performance should not be interpreted as evidence of successful detection of acute stress states. Rather, these findings suggest that the models capture aspects of longer-term stress burden more reliably than transient fluctuations in subjective stress. Moreover temporal aggregation simplifies the prediction task by averaging over short-term fluctuations, such analyses remain informative for several reasons. First, they help determine whether wearable physiology contains information about longer-term stress burden even when momentary stress states cannot be reliably predicted. Second, improvements following aggregation can provide insight into the sources of model error: if performance increases substantially after averaging, this suggests that part of the prediction error arises from short-term variability, temporal misalignment, or measurement noise rather than a complete absence of physiological signal. Finally, aggregated predictions may be relevant for applications in which the goal is not real-time stress detection but the monitoring of sustained stress exposure over days or weeks. Such longer-term indicators may be useful for risk stratification, early detection of prolonged stress burden, and guiding preventive or therapeutic interventions.

To distinguish whether the model captures slow stress buildup or momentary stress dynamics, we decomposed prediction performance into between-week and within-week components. Weeks were defined as consecutive 7-day windows within each participant’s monitoring period rather than calendar weeks. This approach avoids artificial discontinuities at calendar boundaries (e.g., Sunday versus Monday) and ensures that temporally adjacent observations remain grouped within the same aggregation window whenever possible. The between-week (correlation between weekly mean predicted stress and weekly mean reported stress within each user) correlation quantified how well weekly mean predicted stress tracked weekly mean reported stress within each user, whereas the within-week deviation correlation (correlation between deviations from each week’s mean prediction and deviations from each week’s mean reported stress within each user) quantified whether deviations from the weekly mean prediction tracked deviations from the weekly mean reported stress. The model showed stronger performance for between-week variation than for within-week fluctuations (median between-week *r* = 0.27 vs. median within-week *r* = 0.13), indicating that predictions more strongly reflect slower-changing baseline shifts than moment-to-moment stress dynamics.

## V. Discussion

This study aimed to examine whether wearable-derived physiological and motion signals can be used to estimate individuals’ subjective stress states, as assessed by concurrently collected EMA reports, and whether this mapping generalizes across individuals, model classes, and sensing modalities. Our findings are consistent with growing evidence that wearable-based stress detection generalizes poorly across individuals in naturalistic settings [17], [5], [9], [6], [10], [11]. Rather than treating this as a modeling limitation, we show that the failure persists across modalities and model classes, suggesting that the relationship between wearable physiology and subjective stress is fundamentally person-specific.

Feature-based intra-user models consistently outperformed both feature-based inter-user models and end-to-end approaches, indicating that stress physiology is strongly individualized. This supports the view that stress is shaped by person-specific baselines, physiological reactivity, behavioral context, and subjective appraisal (both cognitive and affective), all of which are obscured when data are pooled across individuals. Although autonomic activation is associated with stress, the mapping between physiology and subjective experience appears highly context-dependent, person-specific, and partially non-specific to stress itself. This does not imply that physiological stress detection is impossible, but rather that the search for a single generalized “gold-standard” wearable stress biomarker may be fundamentally limited in ecologically valid settings. The present findings also support theoretical arguments that subjective stress and physiological stress responses are only partially aligned constructs [10], [6], [36].

Higher performance reported in previous wearable stress studies likely reflects differences in task formulation and experimental setting. For example, Gjoreski et al. [14] addressed binary stress-event classification under semi-controlled conditions, whereas we evaluated continuous EMA-based stress detection in fully naturalistic settings using strict LOSO validation. Similarly, Smets et al. [17] focused on population-level physiological associations rather than cross-individual stress detection.

We also observed that the learning depends primarily on exposure to informative stress-related events rather than overall monitoring duration. Models appear to improve when they observe a wider range of stress states, particularly elevated states that provide informative contrast against baseline physiology. Moreover, the prediction errors were highly structured rather than random. Residuals were positively associated with high valence and high arousal, and negatively associated with low valence and low arousal states, showing that errors followed a coherent affective pattern rather than reflecting arbitrary noise. The model captures meaningful stress-related information, but does not fully isolate stress from overlapping affective and physiological processes. Physiological activation can arise from multiple sources, including positive arousal, movement, and contextual demands. Even when the dominant signal aligns with stress, these overlapping processes make perfect construct specificity difficult to achieve [5].

Temporal analyses further clarified that the models seem more sensitive to longer-term buildup or background stress level than to transient changes at the EMA timescale. These findings are consistent with allostatic load, the cumulative physiological effects of repeated or prolonged stress [37]. The findings therefore have important translational implications. Their stronger performance for slower temporal variation, combined with systematic underestimation of high-stress states, suggests that they may be most useful for identifying gradual changes in burden, sustained deviations from baseline, or longer-term risk trajectories rather than detecting acute episodes in real time.

These findings also have implications for foundation models. While large pretrained models may learn general physiological representations, our results suggest that substantial personalization will remain necessary because the physiology– stress relationship is largely individual-specific. Accordingly, we did not evaluate foundation models in this study, as our objective was to characterize the limits of population-level supervised learning rather than investigate pretraining strategies. Consequently, poor inter-user performance should not necessarily be interpreted as evidence against large-scale physiological pretraining, but rather as evidence that successful stress prediction may require a combination of shared representations and user-specific calibration.

### 1) Limitations

Several limitations should be considered. First, participants were monitored for approximately three weeks, limiting exposure to diverse and repeated high-stress events and potentially reducing the stability of personalized models. Second, the cohort consisted of healthy adults from a restricted age range and geographic region, with individuals exhibiting limited stress variability excluded from analyses. Consequently, the findings may not generalize to clinical populations or settings with greater stress diversity, and the relatively low prevalence of intense stress may have hindered learning of high-stress states. Finally, EMA provides an imperfect reference standard. Self-reported stress is subject to temporal imprecision, individual differences in scale use, and reporting variability, meaning that some prediction error likely reflects label noise rather than model failure.

### 2) Conclusions and Future Work

Overall, individualized models consistently outperformed population-level models in detecting subjective stress, but explained only a moderate proportion of its variance. Thus, wearable-derived signals may support scalable, unobtrusive long-term mental health monitoring, yet appear insufficient for robust detection of momentary stress in real-world settings. Practical implementation will likely require personalized calibration, potentially through foundation-model approaches.

Several directions emerge from these findings. First, future work should investigate longer-term personalized monitoring to determine the amount of calibration data required and whether broader exposure to high-stress states improves detection performance. Second, incorporating contextual information (e.g., activity, sleep, environmental factors, and brief self-reports) may improve the specificity of wearable-based stress detection by distinguishing stress from other forms of arousal. Third, modeling approaches that jointly capture broader affective dimensions, such as valence and arousal, may better reflect the information encoded in wearable data than stress alone. Finally, longitudinal studies should evaluate whether personalized wearable-derived signatures predict clinically meaningful outcomes, such as burnout and aggression. Together, these future directions may clarify whether wearable sensing can be used for detecting momentary acute stress, or whether it is more suitable for context-aware monitoring of sustained deviations from an individual’s typical behavioral and physiological state.

## Acknowledgment

Author Peter de Looff was funded by a ZonMw Fellowship grant (06360322210023). For the purpose of Open Access the author has applied a CC BY public copyright licence to any Author Accepted Manuscript version arising from this submission. Author Erno Hermans was supported by a VICI grant from the Dutch Research Council (VI.C.211.106).

