## Supplementary Material for "Detection of Stress in Naturalistic Settings Through Passive Mobile Sensing"

### I. Details of the Preprocessing

*a) Electrodermal activity (EDA):* signals were low-pass filtered (1 Hz Butterworth) to attenuate high-frequency noise [1], [2], automatically screened for artifacts using EDA-Explorer [3], and subjected to continuity/quality checks (minimum 2 minutes segments). Clean data were decomposed into tonic and phasic components using cvxEDA implemented in NeuroKit2 [2], [4]. From these components, standard time- and frequency-domain features were extracted [5], [6] (See Table **??**).

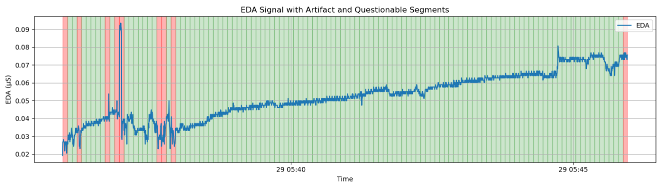


(a)

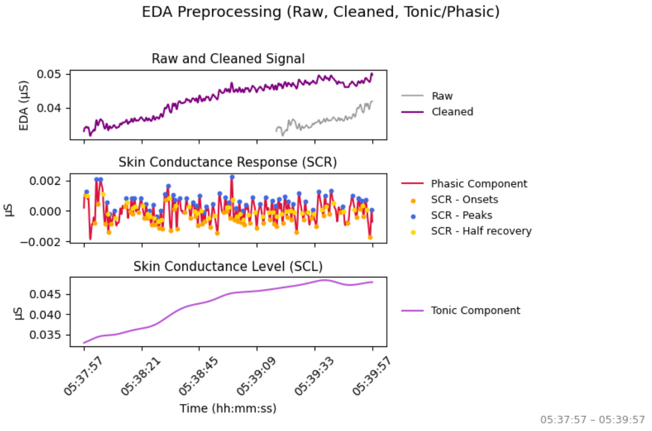


(b)

Fig. 1: EDA preprocessing overview. (a) Full EDA signal showing detected artifacts and retained segments. (b) Zoomed-in 2-min cleaned window with phasic and tonic components and detected SCR peaks, onsets, and recoveries.

*b) Blood Volume Pulse (BVP):* signals were band-pass filtered (0.5–3.3 Hz Butterworth) to isolate cardiac oscillations [7], cleaned using derivative-based quality metrics and reconstruction (pyPPG/E2E-PPG) [8], [9], [10], and peak-detected using the Elgendi method (NeuroKit2) [4], [11]. Inter-beat intervals were derived to compute pulse-rate variability (PRV) features across time, frequency, and nonlinear domains [12], [13] (See Table **??**). Consistent with prior work, variability from BVP is interpreted as PRV rather than HRV due to wearable-specific limitations [14], [15].

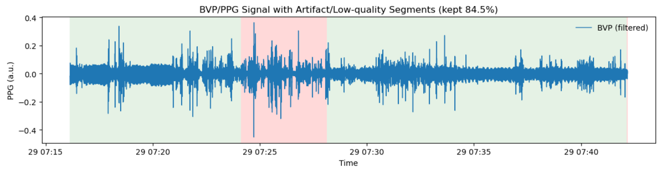


(a)

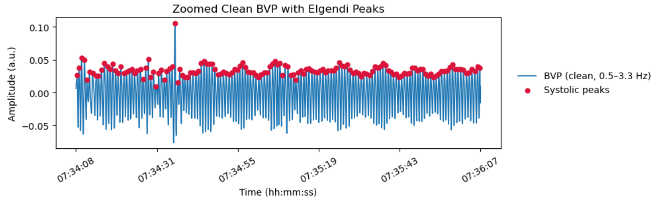


(b)

Fig. 2: BVP signal preprocessing and peak detection. (a) Full wrist PPG/BVP segment showing artifact and low-quality intervals (red) removed; retained clean portions (green) proceed to further analysis. (b) Zoomed 2-min segment with detected peaks used for inter-beat interval and pulse-rate variability analysis.

*c) Accelerometer (ACC):* channels were combined as vector magnitude to quantify overall movement intensity [16], [17]. The magnitude signal was low-pass filtered (10 Hz Butterworth) and median-smoothed (kernel = 3) to remove spikes, based on the Flirt package implementation. Low-variance segments indicative of sensor dropout or non-wear were excluded. Time and frequency-domain features were extracted to quantify movement intensity and detect motion-related artifacts.

*d) Temperature (TEMP):* was low-pass filtered (0.05–0.1 Hz Butterworth) to preserve the slow thermal trends [18] and processed for non-wear detection following DETACH-style criteria [19], [20], [21], [22]: periods with temperature $< 30°$ and $\frac{\Delta T}{\Delta t} < -0.2°$ per min were marked as non-wear, and those $> 26°$ or $\frac{\Delta T}{\Delta t} > 0.1°$ per min as re-wear. Non-wear windows were excluded.

### II. Details of the Model Architectures

This section summarizes the model architectures used in the (i) feature-extracted setting and (ii) raw-data setting. All models were trained as regressors with a single scalar output neuron and linear activation (target: EMA score). When deep learning models are used, regularization and dropout follow the implementation provided in the code.

#### A. Feature-Extracted Models

Feature-extracted inputs were represented as either (a) tabular feature vectors (classical ML, MLP) or (b) short temporal sequences with shape $(T, F)$ for sequence models, where $T$

is the number of timesteps and $F$ the number of features per timestep.

All neural networks were trained for up to 100 epochs with early stopping (patience = 10) based on validation loss. A batch size of 32 was used. The Adam optimizer was employed with an initial learning rate of 0.001. A learning rate scheduler (ReduceLROnPlateau,a factor of 0.1 after 15 epochs without improvement in validation loss) was applied to reduce the learning rate during training based on validation performance.

*a) Classical ML baselines:*

- **Linear Regression (LIN).** Ordinary least squares regression.
- **Ridge Regression (RID).** $\ell_2$-regularized linear regression with $\alpha = 1.0$.
- **Lasso Regression (LAS).** $\ell_1$-regularized regression with $\alpha = 0.1$ and `max_iter` $= 1000$.
- **K-Nearest Neighbors (KNN).** $k = 7$ neighbors, uniform weights, default distance metric (scikit-learn), parallelized with `n_jobs` $= -1$.
- **Random Forest (RF).** 500 trees, maximum depth 30, `min_samples_split` $= 10$, `min_samples_leaf` $= 1$, `max_features` $= None$, bootstrap sampling enabled, out-of-bag scoring enabled, `random_state` $= 42$.
- **XGBoost (XGB).** Gradient-boosted trees with 500 estimators, `max_depth` $= 10$, `learning_rate` $= 0.05$, `subsample` $= 0.8$, `colsample_bytree` $= 0.8$, objective `reg:squarederror`, `random_state` $= 42$.

*b) MLP regressor (tabular):* A feedforward neural network (**MLP**) was implemented in Keras with batch normalization and dropout:

- Dense(256, ReLU) + BatchNorm + Dropout(0.3)
- Dense(128, ReLU) + BatchNorm + Dropout(0.3)
- Dense(64, ReLU) + Dropout(0.3)
- Dense(1, Linear)

The model was compiled with Adam (`lr` $= 0.001$), mean-squared error loss, and MAE as an auxiliary metric.

*c) 1D-CNN regressor (sequence):* A temporal **1D-CNN** was used for sequence inputs $(T, F)$ with $\ell_2$ weight decay ($\lambda = 0.001$), batch normalization, and dropout:

$$\begin{aligned}
&\text{Conv1D}(64, k = 2, \text{ReLU}, \text{same}) \rightarrow \text{BN} \\
&\rightarrow \text{Conv1D}(64, k = 2, \text{ReLU}, \text{same}) \rightarrow \text{BN} \\
&\rightarrow \text{Conv1D}(32, k = 2, \text{ReLU}, \text{same}) \rightarrow \text{BN} \\
&\rightarrow \text{GlobalMaxPool1D} \\
&\rightarrow \text{Dense}(64, \text{ReLU}) \rightarrow \text{Dropout}(0.3) \\
&\rightarrow \text{Dense}(32, \text{ReLU}) \rightarrow \text{Dense}(1, \text{Linear})
\end{aligned}$$

*d) Simple RNN regressor (sequence):* A recurrent baseline (**RNN**) used a single SimpleRNN layer with 64 units and $\ell_2$ regularization ($\lambda = 0.001$):

$$\text{SimpleRNN}(64) \rightarrow \text{Dense}(64) \rightarrow \text{Dropout}(0.3) \rightarrow \text{Dense}(1)$$

*e) BiLSTM regressor (sequence):* A bidirectional LSTM (**BiLSTM**) used a single BiLSTM layer with 64 units and $\ell_2$ regularization ($\lambda = 0.001$):

$$\text{BiLSTM}(64) \rightarrow \text{Dense}(64) \rightarrow \text{Dropout}(0.3) \rightarrow \text{Dense}(1)$$

*1) Raw-Data Models:* Raw inputs were represented as small multi-channel images/tensors (e.g., $32 \times 32 \times C$). All raw-data models were configured as regressors with a final $\text{Dense}(1, \text{Linear})$ output.

*a) Mini-GoogLeNet regressor (Inception-style):* A compact Inception-v1 style network (**GoogLeNet**) was implemented for small inputs. It includes:

- A lightweight stem: Conv2D(64, 3×3, ReLU) + MaxPool
- Two Inception blocks using parallel $1\times1$, $1\times1 \rightarrow 3\times3$, $1\times1 \rightarrow 5\times5$, and pooling$\rightarrow 1\times1$ projections
- Global average pooling, Dense(512, ReLU), Dropout(0.3), Dense(1, Linear)

*b) ResNet-152 regressor (transfer learning):* A **ResNet-152** backbone was used via `tf.keras. applications. ResNet152` with `include_top=False`. Because the raw inputs are $32 \times 32$, inputs were upscaled in-model using `Resizing(resize_to, resize_to)` (default: 224). ImageNet preprocessing was applied (`resnet.preprocess_input`). The head consists of GlobalAveragePooling2D, Dense(512, ReLU), Dropout(0.2), and Dense(1, Linear). When pretrained weights are enabled, the backbone is initially frozen and optionally fine-tuned by unfreezing the last `fine_tune_at` layers (default: last 10 layers). Although recurrence plots differ from natural images, ImageNet pretraining provides a strong initialization by capturing generic visual primitives such as edges, textures, and spatial repetition patterns. These features are directly relevant to recurrence plots, which encode dynamical structure through repeated diagonal lines, texture density, and self-similarity patterns.

Consequently, transfer learning improves optimization stability, accelerates convergence, and acts as an implicit regularizer, particularly under limited physiological datasets. Fine-tuning the final layers further adapts the network to domain-specific recurrence plot characteristics.

*c) AlexNet-style regressor:* A custom **AlexNet**-style CNN was used with multiple Conv2D blocks (kernel size $2\times2$) interleaved with batch normalization and max pooling. The network ends with Flatten and two dense layers:

$$\begin{aligned}
&\text{Dense}(1024, \tanh) \rightarrow \text{Dropout}(0.2) \rightarrow \text{Dense}(256, \tanh) \\
&\rightarrow \text{Dropout}(0.2) \rightarrow \text{Dense}(1)
\end{aligned}$$

This model was used as a regression adaptation of an AlexNet-like design tailored to small input tensors.

*d) DenseNet-style regressor:* A DenseNet-inspired architecture (**DenseNet**) was implemented using dense blocks with bottleneck structure: BN $\rightarrow$ ReLU $\rightarrow$ $1\times1$ Conv (expansion) followed by BN $\rightarrow$ ReLU $\rightarrow$ $3\times3$ Conv (growth), with feature map concatenation. Two dense blocks (repetitions [6,12]) are separated by transition layers that halve channels and downsample via average pooling. The regression head uses GlobalAveragePooling2D and Dense(1, Linear). Default settings included growth rate 48, dropout 0.2 within dense blocks, and $\ell_2$ weight decay $10^{-4}$.

### III. Details of the Recurrence Plot Generation

Recurrence plots (RPs) are a non-linear data visualization technique for exploring the recurrent behavior of dynamical systems in phase space [23]. Given a univariate time series $\{x_i\}$, one reconstructs its trajectory in an $m$-dimensional phase space via time-delay embedding. The RP is then defined as the binary matrix

$$R_{i,j}(\varepsilon) = \Theta\big(\varepsilon - \|\mathbf{x}_i - \mathbf{x}_j\|\big),$$

where $\mathbf{x}_i = [x_i, x_{i+\tau}, \ldots, x_{i+(m-1)\tau}]$, $\varepsilon$ is a threshold, and $\Theta$ is the Heaviside function.

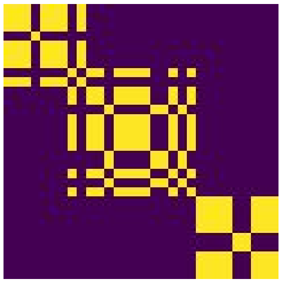

(a) Example 1

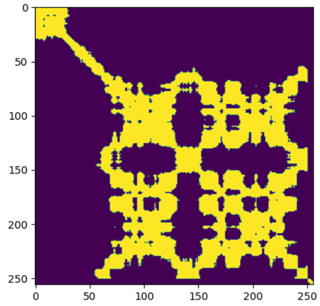


(b) Caption 2

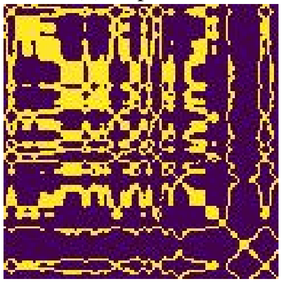

(c) Caption 3

Fig. 3: Examples of recurrence plots generated from our own dataset

The resulting plot reveals diagonal, vertical and horizontal line structures that reflect periodicity, laminar states, and transitions in the underlying dynamics [24].

*a) Benefits for Time-Series Representation:* By converting a one-dimensional signal into a two-dimensional image encoding its recurrence structure, RPs capture both short- and long-term dependencies, nonstationarities, and regime changes in a single, compact representation [25]. Unlike sliding-window statistics or spectrograms, RPs do not assume stationarity or linearity; they sensitively detect transient events, unstable periodic orbits, and subtle dynamical shifts that are often imperceptible in the raw signal or its Fourier spectrum [24].

*b) Superiority over Other Image-Based Techniques:* Compared to other time-series-to-image transforms—such as Gramian Angular Fields [26] or Markov Transition Fields [27]-recurrence plots offer three key advantages:

1) **Dynamical Fidelity:** RPs directly preserve the phase-space geometry of the system, allowing downstream models (e.g. convolutional neural networks) to learn from true dynamical invariants rather than heuristic encodings.
2) **Model-Agnostic Thresholding:** The choice of threshold $\varepsilon$ provides a principled way to tune sensitivity to state changes, enabling multi-scale analysis without redesigning the transform.
3) **Proven Diagnostic Power:** RPs and their quantitative extensions (recurrence quantification analysis) have been widely validated in fields as diverse as cardiology, neurology, and climatology for detecting pathological rhythms, epileptic seizures, and climate regime shifts [28], [25].

### IV. Feature-Extracted Inter-User Models

*a) Model Result Details:* Detailed model-level results for each modality configuration are provided in the supplementary material under `$Appendix/Tables/FeatureExtractedInterUser/Models$`. The corresponding statistical testing results are provided under `$Appendix/Tables/FeatureExtractedInterUser/Statistical\_Testing$`. This appendix summarizes the main performance patterns across modality configurations and reports the aggregate statistical testing results.

Overall (see Tables I and II), the feature-extracted inter-user models showed consistently negative mean $R^2$ values across modality configurations, indicating limited generalization under the inter-user setting. Among the unimodal models, EDA performed best when using MLP, whereas BVP favored Bi-LSTM for error-based metrics. For dual-modality configurations, BVP+EDA favored XGBoost, while EDA+TEMP and BVP+TEMP favored Bi-LSTM. For higher-order modality combinations, BVP+EDA+TEMP favored RNN, whereas the full BVP+EDA+TEMP+ACC configuration favored MLP and XGBoost. Statistical testing did not indicate strong evidence of significant differences across configurations based on the aggregate p-values, although BVP+EDA+TEMP and the full multimodal configuration showed the lowest average p-values. All p-values were $> 0.05$.

Statistical comparisons across models were conducted using LOSO subjects as the paired unit, yielding one subject–level $R^2$ value per model. Given potential non–normality and boundedness of subject–level $R^2$ distributions, the Wilcoxon

TABLE I: Summary of the best-performing feature-extracted inter-user models across modality configurations. Best models were selected separately for each metric based on mean performance. For $R^2$, higher values indicate better performance; for MAE and RMSE, lower values indicate better performance.

| Modality Configuration | Best $R^2$ Model | Best MAE Model | Best RMSE Model |
|---|---|---|---|
| BVP | Lasso / Linear / Ridge | Bi-LSTM | Bi-LSTM |
| EDA | MLP | MLP | MLP |
| BVP+EDA | XGBoost | XGBoost | XGBoost |
| BVP+ACC | Bi-LSTM / Random Forest | Bi-LSTM / Lasso / Linear / Ridge | Lasso |
| BVP+TEMP | Bi-LSTM | Bi-LSTM | Bi-LSTM |
| EDA+ACC | MLP / XGBoost | MLP / XGBoost | MLP / XGBoost |
| EDA+TEMP | Bi-LSTM | Bi-LSTM | Bi-LSTM |
| BVP+EDA+ACC | MLP / Random Forest / XGBoost | Bi-LSTM / Lasso | Lasso |
| BVP+EDA+TEMP | RNN | RNN | RNN |
| BVP+EDA+TEMP+ACC | MLP / XGBoost | MLP / XGBoost | MLP / XGBoost |

TABLE II: Aggregate statistical testing results for feature-extracted inter-user models.

| Modality Configuration | Average p-value | Median p-value |
|---|---|---|
| BVP | 0.0720 | 0.0699 |
| EDA | 0.1370 | 0.1200 |
| BVP+EDA | 0.0844 | 0.0827 |
| BVP+ACC | 0.1340 | 0.1699 |
| BVP+TEMP | 0.1320 | 0.1370 |
| EDA+ACC | 0.1300 | 0.1600 |
| EDA+TEMP | 0.6400 | 0.6600 |
| BVP+EDA+ACC | 0.0920 | 0.9380 |
| BVP+EDA+TEMP | 0.0628 | 0.0620 |
| BVP+EDA+TEMP+ACC | 0.0622 | 0.0676 |

signed-rank test was used as the primary inferential test, while paired $t$–tests were additionally computed as a robustness analysis. To control for multiple comparisons across the 45 pairwise tests, $p$–values were corrected using the Benjamini–Hochberg false discovery rate (FDR) procedure across all pairwise model comparisons. Effect sizes are reported as Cohen's $d$ computed on paired differences (row minus column). Statistical significance was assessed at $\alpha = 0.05$ after FDR correction.

*b) Statistical Testing Summary for Feature-Extracted Inter-User Models:* Full pairwise statistical testing results are provided in the supplementary material under `$Appendix/Tables/FeatureExtractedInterUser/Statistical\_Testing$`. This appendix summarizes the main inferential patterns across modality configurations. Pairwise comparisons were conducted on LOSO-subject $R^2$ scores using Wilcoxon signed-rank tests with Benjamini–Hochberg FDR correction. Paired $t$-tests were additionally inspected as a sensitivity analysis, and Cohen's $d$ was used to quantify paired effect sizes.

Overall (see Table III, statistically significant differences were concentrated around a small number of repeated patterns rather than distributed uniformly across all model pairs. KNN produced the most frequent significant FDR-corrected contrasts, especially in multimodal configurations, with representative adjusted $p$-values below 0.001 and large-to-very-large paired effect sizes ($|d| \approx 1.0$–$2.6$). Linear baselines were also frequently separated from nonlinear models, particularly in BVP, EDA, BVP+EDA, and BVP+ACC. Paired $t$-tests generally supported the same qualitative conclusions but were more conservative for several borderline contrasts, especially KNN versus linear baselines where Wilcoxon-adjusted $p$-values were approximately 0.046 but paired $t$-test adjusted $p$-values were above 0.36.

### V. Raw Inter User Models

*a) Model Result Details::* Detailed model-level results for each raw-data modality configuration are provided in the supplementary material under `$Appendix/Tables/RawIntraUser/Models$`. The corresponding statistical testing results are provided under `$Appendix/Tables/RawIntraUser/Statistical\_Testing$`. This appendix summarizes the main performance trends across modality configurations.

Overall, the raw intra-user models also produced negative mean $R^2$ values across all modality configurations, indicating limited predictive performance despite subject-specific training. AlexNet achieved the strongest results for the unimodal BVP and EDA settings, as well as for EDA+TEMP. In contrast, ResNet was the most consistent architecture across multimodal configurations, yielding the best $R^2$, MAE, and RMSE in most BVP-containing combinations. The lowest aggregate p-values were observed for BVP+EDA+ACC, EDA+TEMP, BVP+ACC, and BVP+TEMP, whereas BVP+EDA+TEMP showed substantially higher aggregate p-values, suggesting weaker evidence of model-level separation in that configuration.

*b) Statistical Testing Details:* Full pairwise statistical testing results are provided in the supplementary material under `Appendix/Tables/RawIntraUser/Statistical\_Testing`. This appendix summarizes the main inferential patterns across raw-data modality configurations. Pairwise comparisons were conducted on LOSO-subject $R^2$ scores using Wilcoxon signed-rank tests with Benjamini–Hochberg FDR correction. Paired $t$-tests were additionally inspected as a sensitivity analysis, and Cohen's $d$ was used to quantify paired effect sizes.

Across raw intra-user models, statistical differences were more architecture- and modality-dependent than in the feature-extracted setting. Unimodal BVP and EDA showed no FDR-corrected significant pairwise differences, despite several moderate effect sizes, suggesting substantial overlap in subject-level performance. In contrast, multimodal settings produced clearer separations, especially BVP+EDA, BVP+TEMP, BVP+EDA+TEMP, and the full BVP+EDA+TEMP+ACC configuration. GoogLeNet and ResNet were most frequently

TABLE III: Summary of statistical testing results for feature-extracted inter-user models. Reported values summarize representative FDR-corrected Wilcoxon $p$-values and paired Cohen's $d$ effect sizes.

| Modality Configuration | Main FDR-Corrected Wilcoxon Pattern | Effect Size Pattern | Wilcoxon vs. Paired $t$-test |
|---|---|---|---|
| BVP | Significant contrasts included RF/XGB/MLP vs. linear baselines ($p_{\mathrm{FDR}} = 0.002$–$0.013$) and KNN vs. most models ($p_{\mathrm{FDR}} < 0.001$; KNN vs. RNN $p = 0.002$). | KNN-related effects were large ($\|d\| = 1.10$–$1.52$). Linear vs. nonlinear contrasts were also large ($\|d\| = 0.97$–$1.18$). | Broadly consistent; only minor borderline differences were observed. |
| EDA | Linear models were frequently separated from nonlinear models ($p_{\mathrm{FDR}} < 0.001$). RF differed from XGB/MLP ($p = 0.009$) and KNN ($p = 0.016$). | Large effects appeared for nonlinear vs. linear contrasts, especially KNN vs. LIN/RID/LAS ($d = 2.41$–$2.59$) and XGB/MLP vs. linear models ($\|d\| = 1.31$–$1.38$). | Mostly consistent; RF vs. BiLSTM and LIN vs. RID were Wilcoxon-significant but not paired-$t$ significant. |
| BVP+EDA | RF/XGB/MLP significantly differed from linear baselines ($p = 0.006$–$0.009$). KNN showed broad significant separation from other models ($p_{\mathrm{FDR}} < 0.001$; KNN vs. RNN $p = 0.006$). | KNN-related effects were large ($d = 1.56$–$1.78$). Effects involving CNN were smaller ($d \approx 0.20$–$0.44$). | Broadly consistent, with borderline cases such as linear baselines vs. RF. |
| BVP+ACC | RF/XGB/MLP differed from linear baselines and KNN ($p_{\mathrm{FDR}} < 0.001$). KNN also differed from RNN ($p = 0.009$) and BiLSTM ($p < 0.001$). | KNN effects were very large ($d = 1.76$–$2.22$). Linear models showed large negative effects relative to nonlinear baselines ($\|d\| = 1.26$–$1.29$). | Mostly consistent; LIN/RID vs. LAS were significant under Wilcoxon ($p = 0.020$–$0.024$) but not paired $t$-tests. |
| BVP+TEMP | KNN was the primary significant model, differing from RF/XGB/MLP ($p_{\mathrm{FDR}} < 0.001$), linear baselines ($p = 0.046$), RNN ($p < 0.001$), and BiLSTM ($p < 0.001$). | KNN-related effects were very large against RF/XGB/MLP ($d = 1.87$–$1.90$) and recurrent models ($\|d\| = 2.08$–$2.28$). | KNN vs. LIN/RID/LAS was significant only under Wilcoxon ($p = 0.046$ vs. paired-$t$ $p = 0.42$–$0.45$). |
| EDA+ACC | KNN differed from most models ($p = 0.001$–$0.003$), except CNN. RF also differed from LIN/RID/LAS ($p = 0.027$–$0.050$), RNN ($p = 0.011$), and BiLSTM ($p = 0.015$). | KNN effects were large-to-very-large ($d = 1.57$–$1.99$). Recurrent-model effects were smaller ($d \approx 0.29$–$0.65$). | Several Wilcoxon-only contrasts appeared, including RNN vs. RF and CNN vs. RNN. |
| EDA+TEMP | This setting produced the widest spread of significant comparisons, including RF vs. XGB/MLP ($p = 0.023$), RF vs. KNN ($p = 0.013$), KNN vs. XGB/MLP/RNN/BiLSTM ($p = 0.003$), and KNN vs. linear baselines ($p = 0.023$). | KNN showed large effects against RF/XGB/MLP ($d = 1.34$–$1.83$). Other contrasts were mostly small-to-moderate. | Paired $t$-tests were more conservative; Wilcoxon detected additional significant contrasts such as RF vs. XGB/MLP and KNN vs. LIN. |
| BVP+EDA+ACC | KNN was the only model with broad significant separation, differing from RF/XGB/MLP/LIN/RID/LAS ($p_{\mathrm{FDR}} < 0.001$), RNN ($p = 0.011$), and BiLSTM ($p < 0.001$). | KNN-related effects were large-to-very-large ($d = 1.54$–$1.78$). Most non-KNN effects were small-to-moderate. | No substantive decision reversals were observed between Wilcoxon and paired $t$-tests. |
| BVP+EDA+TEMP | KNN differed from RF/XGB/MLP ($p_{\mathrm{FDR}} < 0.001$), linear baselines ($p = 0.046$), RNN ($p < 0.001$), and BiLSTM ($p < 0.001$). | KNN-related effects were very large against RF/XGB/MLP ($d = 1.93$–$1.99$) and recurrent models ($\|d\| = 1.27$–$1.61$). | KNN vs. LIN/RID/LAS was significant under Wilcoxon ($p = 0.046$) but not paired $t$-tests ($p = 0.415$–$0.417$). |
| BVP+EDA+ACC+TEMP | KNN again showed the clearest separations: KNN vs. RF/XGB/MLP ($p_{\mathrm{FDR}} < 0.001$), vs. LIN/RID/LAS ($p = 0.046$), vs. RNN ($p = 0.022$), and vs. BiLSTM ($p = 0.046$). | KNN-related effects were large against RF/XGB/MLP ($d = 1.64$–$1.75$), RNN ($\|d\| = 1.00$), and BiLSTM ($\|d\| = 0.85$). | KNN vs. LIN/RID/LAS was significant only under Wilcoxon ($p = 0.046$ vs. paired-$t$ $p = 0.365$). |

TABLE IV: Summary of best-performing raw intra-user models across modality configurations. Best models were selected separately for each metric based on mean performance. For R$^2$, higher values indicate better performance; for MAE and RMSE, lower values indicate better performance.

| Modality Configuration | Best R$^2$ Model | Best MAE Model | Best RMSE Model |
|---|---|---|---|
| BVP | AlexNet ($R^2 = -0.4050$) | AlexNet (MAE = 1.1674) | AlexNet (RMSE = 1.4173) |
| EDA | AlexNet ($R^2 = -0.3110$) | AlexNet (MAE = 1.4040) | AlexNet (RMSE = 1.7174) |
| BVP+EDA | ResNet ($R^2 = -0.7761$) | ResNet (MAE = 1.2787) | GoogleNet (RMSE = 1.0413) |
| BVP+ACC | ResNet ($R^2 = -0.6364$) | ResNet (MAE = 1.3322) | ResNet (RMSE = 1.5553) |
| BVP+TEMP | ResNet ($R^2 = -0.4126$) | ResNet (MAE = 1.6949) | ResNet (RMSE = 1.8612) |
| EDA+ACC | ResNet ($R^2 = -0.6244$) | ResNet (MAE = 1.2176) | ResNet (RMSE = 1.4528) |
| EDA+TEMP | AlexNet ($R^2 = -0.2353$) | AlexNet (MAE = 0.9585) | AlexNet (RMSE = 1.3491) |
| BVP+EDA+ACC | ResNet ($R^2 = -0.4967$) | ResNet (MAE = 1.3371) | ResNet (RMSE = 1.5967) |
| BVP+EDA+TEMP | ResNet ($R^2 = -0.3122$) | ResNet (MAE = 1.3688) | ResNet (RMSE = 1.6115) |
| BVP+EDA+TEMP+ACC | ResNet ($R^2 = -0.6707$) | ResNet (MAE = 1.4098) | ResNet (RMSE = 1.6457) |

TABLE V: Aggregate statistical testing results for raw intra-user models.

| Modality Configuration | Average p-value | Median p-value |
|---|---|---|
| BVP | 0.1300 | 0.1350 |
| EDA | 0.1400 | 0.1400 |
| BVP+EDA | 0.1340 | 0.1350 |
| BVP+ACC | 0.0627 | 0.0640 |
| BVP+TEMP | 0.0640 | 0.0645 |
| EDA+ACC | 0.1300 | 0.1280 |
| EDA+TEMP | 0.0620 | 0.0630 |
| BVP+EDA+ACC | 0.0572 | 0.0576 |
| BVP+EDA+TEMP | 0.5350 | 0.5360 |
| BVP+EDA+TEMP+ACC | 0.0726 | 0.0723 |

involved in significant contrasts, with several large paired effect sizes ($|d| \approx 1.0$–$1.46$). Paired $t$-tests generally supported the same qualitative conclusions, but were more conservative in EDA+ACC and EDA+TEMP, where Wilcoxon-significant contrasts did not remain significant under the parametric sensitivity analysis.

### VI. Feature Extracted Intra User Models

*a) Model Result Details::* Detailed model-level results for each intra-user modality configuration are provided in the supplementary material under `$Appendix/Tables/FeatureExtractedIntraUser/Models$`. This

TABLE VI: Summary of statistical testing results for raw intra-user models. Reported values summarize representative FDR-corrected Wilcoxon $p$-values and paired Cohen's $d$ effect sizes.

| Modality Configuration | Main FDR-Corrected Wilcoxon Pattern | Effect Size Pattern | Wilcoxon vs. Paired $t$-test |
|---|---|---|---|
| BVP | No pairwise comparison remained significant after FDR correction. | Effects were small-to-moderate, with the largest contrasts involving DenseNet vs. AlexNet ($d = 0.66$), DenseNet vs. ResNet ($d = -0.65$), and GoogLeNet vs. DenseNet ($d = -0.73$). | Fully consistent; no comparison was significant after correction under either test. |
| EDA | No pairwise comparison remained significant after FDR correction. | Moderate effects appeared for DenseNet-related contrasts, including AlexNet vs. DenseNet ($d = 0.75$) and DenseNet vs. ResNet ($d = -0.78$). | Fully consistent; no comparison was significant after correction under either test. |
| BVP+EDA | Significant contrasts included AlexNet vs. DenseNet ($p = 0.0013$), AlexNet vs. ResNet ($p = 0.0023$), AlexNet vs. GoogLeNet ($p < 0.0001$), and ResNet vs. GoogLeNet ($p < 0.0001$). | Large effects were observed for GoogLeNet vs. AlexNet ($d = 1.07$), GoogLeNet vs. ResNet ($d = 1.09$), and ResNet vs. AlexNet ($d = -0.814$). | Consistent; all listed Wilcoxon-significant contrasts also remained significant under paired $t$-tests. |
| BVP+ACC | Significant contrasts included AlexNet vs. ResNet ($p = 0.008$), AlexNet vs. GoogLeNet ($p = 0.006$), DenseNet vs. ResNet ($p = 0.021$), and ResNet vs. GoogLeNet ($p = 0.006$). | Effects were large for GoogLeNet vs. AlexNet ($d = 0.924$) and GoogLeNet vs. ResNet ($d = 0.952$), but negligible for ResNet-related contrasts reported as $d = 0.00$. | Mostly consistent; ResNet vs. DenseNet was significant under Wilcoxon but not paired $t$ ($p = 0.181$). |
| BVP+TEMP | Significant contrasts included AlexNet vs. ResNet ($p = 0.0088$), AlexNet vs. GoogLeNet ($p = 0.0078$), DenseNet vs. ResNet ($p = 0.0117$), DenseNet vs. GoogLeNet ($p = 0.0059$), and ResNet vs. GoogLeNet ($p = 0.0059$). | Large effects supported the hierarchy, including GoogLeNet vs. AlexNet ($d = 1.07$), GoogLeNet vs. DenseNet ($d = 1.16$), GoogLeNet vs. ResNet ($d = 1.17$), and ResNet vs. AlexNet ($d = -1.18$). | Highly consistent; all Wilcoxon-significant contrasts also remained significant under paired $t$-tests. |
| EDA+ACC | Significant contrasts included AlexNet vs. DenseNet ($p = 0.029$), AlexNet vs. GoogLeNet ($p = 0.006$), DenseNet vs. ResNet ($p = 0.027$), and ResNet vs. GoogLeNet ($p = 0.006$). | Moderate-to-large effects appeared for DenseNet vs. AlexNet ($d = 0.794$), GoogLeNet vs. AlexNet ($d = 0.751$), and GoogLeNet vs. ResNet ($d = 0.759$). | Paired $t$-tests were more conservative; several Wilcoxon-significant contrasts became non-significant under paired $t$ ($p = 0.062$). |
| EDA+TEMP | Significant contrasts included AlexNet vs. DenseNet ($p = 0.012$), AlexNet vs. GoogLeNet ($p = 0.012$), DenseNet vs. ResNet ($p = 0.012$), and ResNet vs. GoogLeNet ($p = 0.012$). | Moderate effects were observed, especially GoogLeNet vs. AlexNet/ResNet ($d = 0.83$), DenseNet vs. AlexNet ($d = 0.71$), and DenseNet vs. ResNet ($d = -0.71$). | Paired $t$-tests were more conservative; Wilcoxon-significant contrasts were not significant under paired $t$ ($p = 0.128$). |
| BVP+EDA+ACC | Significant contrasts included AlexNet vs. ResNet ($p = 0.005$), AlexNet vs. GoogLeNet ($p = 0.007$), DenseNet vs. GoogLeNet ($p = 0.005$), and ResNet vs. GoogLeNet ($p = 0.005$). | Effects were mostly moderate, including ResNet vs. AlexNet ($d = -0.65$), GoogLeNet vs. AlexNet ($d = -0.71$), GoogLeNet vs. DenseNet ($d = -0.64$), and GoogLeNet vs. ResNet ($d = -0.63$). | Mixed; paired $t$ detected ResNet vs. DenseNet ($p = 0.009$) where Wilcoxon did not. |
| BVP+EDA+TEMP | Significant contrasts included AlexNet vs. ResNet ($p = 0.004$), AlexNet vs. GoogLeNet ($p = 0.004$), DenseNet vs. ResNet ($p = 0.006$), and ResNet vs. GoogLeNet ($p = 0.004$). | Large effects appeared for ResNet vs. AlexNet ($d = -1.461$), GoogLeNet vs. AlexNet ($d = 1.106$), and GoogLeNet vs. ResNet ($d = 1.131$). | Mostly consistent; paired $t$-tests were less sensitive for DenseNet-related moderate effects. |
| BVP+EDA+TEMP+ACC | Significant contrasts included AlexNet vs. ResNet ($p = 0.005$), AlexNet vs. GoogLeNet ($p = 0.012$), DenseNet vs. ResNet ($p = 0.046$), and ResNet vs. GoogLeNet ($p = 0.003$). | Large effects appeared for GoogLeNet vs. AlexNet ($d = 1.317$), GoogLeNet vs. ResNet ($d = 1.320$), and ResNet vs. AlexNet ($d = -1.054$). | Mostly consistent; DenseNet-related contrasts were borderline or non-significant under paired $t$-tests. |

appendix summarizes the main performance trends across modality configurations.

Overall, feature-extracted intra-user models showed stronger performance than the inter-user setting, with several modality configurations achieving positive mean $R^2$ values. XGBoost was the most consistent model across multimodal settings, yielding the best MAE and RMSE in most configurations and the best mean $R^2$ for BVP+EDA+TEMP and BVP+EDA+TEMP+ACC. Random Forest also performed competitively, particularly for BVP+TEMP, while recurrent models such as Bi-LSTM and RNN were more competitive in selected unimodal or EDA-containing settings. The strongest overall configuration was the full BVP+EDA+TEMP+ACC setting, where XGBoost achieved the highest mean $R^2$ (0.32), lowest MAE (1.45), and lowest RMSE (1.68).

#### A. Statistical Testing Summary for Feature-Extracted Intra-User Models

Full pairwise statistical testing results are provided in the supplementary material under `Appendix/Tables/FeatureExtractedIntraUser/Statistical\_Testing`. This appendix summarizes the main inferential patterns across modality configurations. Pairwise comparisons were conducted on LOSO-subject $R^2$ scores using Wilcoxon signed-rank tests with Benjamini–Hochberg FDR correction. Paired $t$-tests were additionally inspected as a sensitivity analysis, and Cohen's $d$ was used to quantify paired effect sizes.

Overall, the feature-extracted intra-user statistical tests showed stronger model separation than the inter-user analyses, particularly for BVP, BVP+EDA, BVP+ACC, EDA+ACC, and EDA+TEMP. Significant differences were most often associated with contrasts between nonlinear models and linear baselines, as well as with XGBoost, Random Forest, KNN, and recurrent models depending on the modality configuration. EDA+TEMP showed the densest pattern of significant pairwise differences, with many adjusted Wilcoxon $p$-values below 0.001. However, higher-order multimodal configurations such as BVP+EDA+ACC, BVP+EDA+TEMP, and BVP+EDA+TEMP+ACC showed weaker Wilcoxon evidence after FDR correction despite large effect-size estimates. In these cases, discrepancies between Wilcoxon and paired $t$-tests suggest that some apparent differences may be sensitive to distributional assumptions or subject-level variability.

TABLE VII: Summary of best-performing feature-extracted intra-user models across modality configurations. Best models were selected separately for each metric based on mean performance. For $R^2$, higher values indicate better performance; for MAE and RMSE, lower values indicate better performance.

| Modality Configuration | Best $R^2$ Model | Best MAE Model | Best RMSE Model |
|---|---|---|---|
| BVP | MLP ($R^2 = -0.10$) | 1D-CNN / RF / XGBoost (MAE = 1.86) | MLP (RMSE = 2.11) |
| EDA | Bi-LSTM ($R^2 = -0.12$) | Bi-LSTM / RNN (MAE = 1.78) | Bi-LSTM / RNN (RMSE = 1.92) |
| BVP+EDA | 1D-CNN ($R^2 = 0.15$) | XGBoost (MAE = 1.50) | XGBoost (RMSE = 1.76) |
| BVP+ACC | Bi-LSTM ($R^2 = 0.09$) | XGBoost (MAE = 1.76) | XGBoost (RMSE = 1.97) |
| BVP+TEMP | Random Forest ($R^2 = 0.11$) | XGBoost (MAE = 1.57) | XGBoost (RMSE = 1.86) |
| EDA+ACC | XGBoost ($R^2 = 0.09$) | XGBoost (MAE = 1.73) | XGBoost (RMSE = 1.96) |
| EDA+TEMP | Bi-LSTM ($R^2 = 0.11$) | XGBoost (MAE = 1.30) | XGBoost (RMSE = 1.53) |
| BVP+EDA+ACC | 1D-CNN ($R^2 = 0.25$) | XGBoost (MAE = 1.49) | XGBoost (RMSE = 1.72) |
| BVP+EDA+TEMP | XGBoost ($R^2 = 0.31$) | XGBoost (MAE = 1.46) | XGBoost (RMSE = 1.72) |
| BVP+EDA+TEMP+ACC | XGBoost ($R^2 = 0.32$) | XGBoost (MAE = 1.45) | XGBoost (RMSE = 1.68) |

TABLE VIII: Aggregate statistical testing results for feature-extracted intra-user models.

| Modality Configuration | Average p-value | Median p-value |
|---|---|---|
| BVP | 0.0613 | 0.0610 |
| EDA | 0.0530 | 0.0560 |
| BVP+EDA | 0.0449 | 0.0436 |
| BVP+ACC | 0.0321 | 0.0436 |
| BVP+TEMP | 0.0592 | 0.0476 |
| EDA+ACC | 0.0687 | 0.0476 |
| EDA+TEMP | 0.0491 | 0.0496 |
| BVP+EDA+ACC | 0.0445 | 0.0462 |
| BVP+EDA+TEMP | 0.0482 | 0.0510 |
| BVP+EDA+TEMP+ACC | 0.0442 | 0.0476 |

[28] J. P. Zbilut and C. L. Webber, "Embeddings and delays as derived from

TABLE IX: Summary of statistical testing results for feature-extracted intra-user models. Reported values summarize representative FDR-corrected Wilcoxon $p$-values and paired Cohen's $d$ effect sizes.

| **Modality Configuration** | **Main FDR-Corrected Wilcoxon Pattern** | **Effect Size Pattern** | **Wilcoxon vs. Paired $t$-test** |
|---|---|---|---|
| BVP | Significant contrasts involved RF/XGB/MLP versus linear baselines and KNN ($p < 0.01$–$0.04$), with KNN differing from CNN/RNN/BiLSTM ($p < 0.01$). | Large effects appeared for KNN vs. RF/XGB ($d = 1.93$–$1.99$) and RNN/BiLSTM vs. linear baselines ($\|d\| \approx 0.92$–$1.00$). | Mostly consistent; Wilcoxon-only differences appeared for RF vs. CNN, MLP vs. CNN, and LIN vs. RID. |
| EDA | RNN and BiLSTM differed from most classical baselines ($p = 0.002$–$0.03$). KNN also differed from several tree/linear models ($p = 0.01$–$0.05$). | RNN/BiLSTM showed large effects against most classical models ($d \approx 0.67$–$0.96$). KNN showed moderate negative effects relative to stronger models. | Highly consistent; only borderline contrasts such as RID vs. KNN differed slightly. |
| BVP+EDA | XGBoost differed from all other models ($p < 0.01$). RF differed from most models except CNN; recurrent models differed from linear and KNN models. | XGBoost showed large effects over RF/MLP ($d = 1.15$ and $1.35$ in magnitude). KNN showed large negative effects versus tree/neural models ($\|d\| > 1.8$). | Highly consistent; disagreements were limited to small-effect contrasts. |
| BVP+ACC | XGBoost and RF showed broad significant separation from linear baselines and KNN ($p < 0.001$–$0.02$). KNN differed from CNN/RNN/BiLSTM ($p < 0.001$–$0.01$). | KNN-related effects were very large ($d = 1.84$–$2.72$). XGBoost and RF also showed large effects against linear/MLP comparisons. | Mostly consistent, with only threshold-level differences such as XGB vs. BiLSTM. |
| BVP+TEMP | RF/XGBoost differed from linear baselines and KNN ($p = 0.01$–$0.02$). CNN differed from linear baselines ($p = 0.02$–$0.03$). | XGBoost-related effects were large ($\|d\| \approx 1.3$–$2.0$). CNN/RNN/BiLSTM showed moderate-to-large effects versus linear baselines. | Largely consistent; moderate-effect RF vs. RNN/BiLSTM remained non-significant under both tests. |
| EDA+ACC | RF/XGBoost/MLP and KNN differed strongly from linear baselines ($p < 0.001$–$0.05$). CNN/RNN/BiLSTM also differed from linear baselines. | KNN-related effects were very large versus RF/XGB/MLP ($d = 1.43$–$3.15$). CNN/RNN/BiLSTM showed large effects versus linear baselines. | Broadly consistent; KNN vs. LAS was borderline under Wilcoxon ($p = 0.05$) but not paired $t$. |
| EDA+TEMP | This configuration showed the densest separation pattern. RF and XGBoost differed from nearly all models ($p = 0.00026$–$0.00086$). CNN also differed from several classical baselines ($p = 0.0054$–$0.0063$). | Very large effects appeared for RF/XGBoost versus MLP/KNN ($\|d\| \approx 1.85$–$2.10$). CNN showed moderate effects versus linear/KNN models ($d \approx 0.65$–$0.72$). | Mostly consistent; minor disagreements occurred near threshold, such as MLP vs. LIN. |
| BVP+EDA+ACC | No pairwise comparison remained significant after BH-FDR correction. | Despite non-significance, several effects were moderate-to-large ($\|d\| \approx 1.0$–$1.5$), suggesting unstable subject-wise differences. | Fully aligned; all comparisons remained non-significant under both Wilcoxon and paired $t$-tests. |
| BVP+EDA+TEMP | No Wilcoxon comparison remained significant after BH-FDR correction. | Some effect-size estimates were large, including KNN vs. RF/XGB ($d > 1$), suggesting instability or outlier sensitivity. | Paired $t$-tests detected several significant contrasts, whereas Wilcoxon did not, indicating sensitivity to distributional assumptions. |
| BVP+EDA+TEMP+ACC | No Wilcoxon comparison remained significant after BH-FDR correction. | Effects ranged from moderate to very large, including LAS/KNN vs. RF/MLP ($d \approx 2.5$) and RNN vs. BiLSTM ($d = -2.5$). | Paired $t$-tests identified several significant contrasts, but Wilcoxon remained fully non-significant, supporting the more conservative interpretation. |

quantification of recurrence plots," *Physics Letters A*, vol. 171, no. 3–4, pp. 199–203, 1992.
